# The Shape of Trees – Limits of Current Diversification Models

**DOI:** 10.1101/2021.01.26.428344

**Authors:** Orlando Schwery, Brian C. O’Meara

## Abstract

To investigate how biodiversity arose, the field of macroevolution largely relies on model-based approaches to estimate rates of diversification and what factors influence them. The number of available models is rising steadily, facilitating the modeling of an increasing number of possible diversification dynamics, and multiple hypotheses relating to what fueled or stifled lineage accumulation within groups of organisms. However, growing concerns about unchecked biases and limitations in the employed models suggest the need for rigorous validation of methods used to infer. Here, we address two points: the practical use of model adequacy testing, and what model adequacy can tell us about the overall state of diversification models. Using a large set of empirical phylogenies, and a new approach to test models using aspects of tree shape, we test how a set of staple models performs with regards to adequacy. Patterns of adequacy are described across trees and models and causes for inadequacy – particularly if all models are inadequate – are explored. The findings make clear that overall, only few empirical phylogenies cannot be described by at least one model. However, finding that the best fitting of a set of models might not necessarily be adequate makes clear that adequacy testing should become a step in the standard procedures for diversification studies.

## Introduction

Among the main goals of the study of macroevolution is to explain large scale patterns of biodiversity, and to unveil the mechanisms behind it. In the case of the diversity of species, and particularly the heterogeneity of how that diversity is distributed both in space and across the tree of life, model-based diversification studies have become the method of choice. Since the use of models explicitly allows the inclusion of particular mechanisms and influencing factors into the analyses, making them particularly useful for testing specific hypotheses, rather than just describing patterns. Therefore, such models are especially promising to not only explain diversity patterns, but also better understand the forces that generate this diversity. This has led to considerable development of new approaches, ranging from simple estimation of constant speciation rates (λ) and extinction rates (μ), to via rates influenced by time or diversity, to more complex sets of models in which diversification rates depend on trait states (Maddison, Midford & Otto 2007; FitzJohn, Maddison & Otto 2009; FitzJohn 2010; FitzJohn 2012) or geographic areas (Goldberg, Lancaster & Ree 2011), or approaches that focused on the localization of shifts in diversification rates (Alfaro *et al.* 2009; Rabosky 2014), all of which are widely used.

However, all of these approaches have underlying assumptions, which may not always be satisfied by empirical data; and they all have limitations, which can severely bias our results and mislead our interpretations thereof. Examples for this include biases introduced by under-parametrized substitution models (Revell, Harmon & Glor 2005), a lack of replication for categorical traits which only occur in one clade (Maddison & FitzJohn 2015), models erroneously inferring trait effects on diversification rates when the trait is actually neutral (Rabosky & Goldberg 2015), and the fact that most studies only rely on the explanatory power of a model to assess its adequacy, which is often not sufficient (Pennell et al. 2015). While ways to address some of these issues through more elaborate models (Beaulieu & O’Meara 2016; Caetano, O’Meara & Beaulieu 2018) or explicit adequacy testing (Höhna, May & Moore 2015) have been proposed, a widespread standardized use of model adequacy testing has not yet established in the field (Brown & ElDabaje 2008), suggesting that the magnitude of model adequacy problems in the field of diversification studies is not yet known.

Here, we employ a simulation-based approach using tree shape metrics to assess the adequacy of standard diversification models across a large set of empirical trees. The method is implemented in the R package *BoskR*, which is described in detail in Schwery and O’Meara (2020). The results of the assessment should show us how well the models perform overall in comparison and reveal patterns on which aspects of the tree shapes they might fail to describe. However, trees for which all models fail might be indicative of common weaknesses of all current models and exploring the causes of inadequacy might bring to light properties of trees for which none of our current models can account. Exploring these could allow the identification of the kind of models we might still be missing and where the field has to develop further.

## Material and Methods

### Empirical Phylogenies

We assembled a collection of empirical trees of various sizes and taxonomic groups. Using the R package *datelife* (Stoltzfus *et al.* 2013; Nguyen *et al.* 2018), we assembled all chronograms from the Open Tree of Life (Hinchliff et al. 2015) that had between 30 and 200 taxa. We added the widely used tree of 87 whale species (Steeman et al. 2009), including its subtrees of the clades Balaenopteridae, Phocoenidae, and Phyllostomidae, and a tree of 11 species of *Calomys* (Pigot, Owens & Orme 2012). Finally, we added trees of 77 and 470 species respectively of Ericaceae (derived from Schwery et al. 2015), 575 species of Poales (Bouchenak-Khelladi, Muasya & Linder 2014), 584 species of Fagales (Xing *et al.* 2014), 309 species of Liverworts (Laenen *et al.* 2014), and 549 species of Angiosperms (O’Meara *et al.* 2016) coming to a total of 131 trees (the list of which is given in Table S1).

Additionally, we obtained a tree set of 214 ultrametric trees of vertebrate families, which was initially used by Lewitus and Morlon (2016b) to analyze patterns of diversification with their new method using Laplacian spectra (Lewitus & Morlon 2016a), which we are making use of here. The same set was subsequently used by Burin *et al.* (2019) to investigate how well one can estimate diversification rates under scenarios of diversity decline.

Combined, our set consisted of 345 trees spanning a broad range of taxonomic groups. Before subsequent analyses, we checked whether the trees were ultrametric and strictly bifurcating, and corrected both where necessary using *BoskR*’s *TreeCorr* function.

### Adequacy Test

From the diversification models available in *BoskR*, we chose a set of six basic models to represent some of the different rate dynamics that are implemented. Those were 1) constant speciation and no extinction (Yule), 2) constant speciation and extinction, time-dependent birth death with, 3) exponential speciation and constant extinction, 4) constant speciation and exponential extinction, as well as diversity-dependent birth death with, 5) linear speciation and constant extinction, and 6) linear speciation and extinction.

To determine the adequacy of our candidate models to describe the diversification dynamics behind our empirical tree sets, we made use of the R package *BoskR* (Schwery and O’Meara 2020). For each tree, we estimated diversification rates (and additional model parameters in case of time- and diversity-dependent models) under each model, and then used those estimates to simulate 1000 trees per model and tree. We then calculated the three shape metrics (principal Eigenvalue, skewness, and peakedness) from their spectral density profiles, and compared the metrics of each empirical tree with its corresponding simulated trees. Model inadequacy was determined both via Bonferroni-corrected p-values and 2D convex-hulls, with inadequate models showing significant p-values (<0.05) or having the empirical tree’s metrics lie outside of at least one of the three polygons.

### Trees without adequate Models

When all employed models are inadequate for a tree, this could mean that we have yet to develop suitable models to describe its underlying mode of diversification. While testing all available models and model-variations currently in existence is beyond the scope of this work, we wanted to expand the basic model set used by eleven, to get a sense of how many trees could actually not be accounted for by any model. These additional models were time dependent birth death with: 7) constant speciation and linear extinction, 8) exponential speciation and extinction, 9) exponential speciation and no extinction, 10) exponential speciation and linear extinction, 11) linear speciation and constant extinction, 12) linear speciation and extinction, 13) linear speciation and exponential extinction, 14) linear speciation and no extinction, as well as diversity-dependent birth death with: 15) exponential speciation and constant extinction, 16) constant speciation and linear extinction, 17) constant speciation and exponential extinction. We tested the adequacy of these models on the trees for which all previous models were inadequate, using the same procedure as described above.

For any trees for which still no model was adequate, we investigated the causes of inadequacy of the models by comparing the tree’s metrics to those of the others, and by more closely inspecting its spectral density, and taxonomical background.

### Adequacy and Fit

An important consideration is whether model adequacy is related to model fit, that is, whether adequate models tend to fit the data well, whereas inadequate models do not, or, in other words, whether mismatches are possible. To that end, we compared the fits of the different models by comparing their respective AIC. We first obtained the log-likelihood associated to parameter estimates under each model. Because different R packages calculate the likelihood differently, those values are not directly comparable (Stadler 2013). While we could transform the likelihoods of the models that were implemented in *ape* (Yule, constant-rate birth-death), to match those from *DDD* (the two diversity-dependent models) by dividing them by −2, the models implemented in *RPANDA* (the two time-dependent models) used a different likelihood equation altogether, which made it impossible to compare them by simple transformations (see Table 1 in Stadler 2013). We thus excluded those two models from the comparison. For the remaining four models, we calculated the AIC from the adjusted log-likelihoods, determined the best model using ΔAIC, and tallied how often the best fitting model and the remaining models were either adequate or inadequate.

**Table 1:** Model Inadequacy by Model. Numbers and percentages of trees for which a particular model was adequate or inadequate for, assessed using either Bonferroni-corrected p-values (above) or 2D convex-hulls (below). BD=constant rate birth-death, T_ex_co=time-dependent birth-death with exponential λ and constant μ, T_co_ex=time-dependent birth-death with constant λ and exponential μ. DD_lin_co=diversity-dependent birth-death with linear λ and constant μ. DD_lin_lin=diversity-dependent birth-death with linear λ and linear μ.

| Model | Yule | BD | T_ex_co | T_co_ex | DD_lin_co | DD_lin_lin |
| --- | --- | --- | --- | --- | --- | --- |
| **Bonferroni-Pvalues** |  |  |  |  |  |  |
| **Adequate** | 201 | 209 | 199 | 193 | 201 | 171 |
| **Inadequate** | 33 | 25 | 35 | 41 | 33 | 63 |
| **% adequate** | 85.897 | 89.316 | 85.043 | 82.479 | 85.897 | 73.077 |
| **% inadequate** | 14.103 | 10.684 | 14.957 | 17.521 | 14.103 | 26.923 |
| **2D convex-hulls** |  |  |  |  |  |  |
| **Adequate** | 201 | 205 | 183 | 185 | 203 | 187 |
| **Inadequate** | 33 | 29 | 51 | 49 | 31 | 47 |
| **% adequate** | 85.897 | 87.607 | 78.205 | 79.060 | 86.752 | 79.915 |
| **% inadequate** | 14.103 | 12.393 | 21.795 | 20.940 | 13.248 | 20.085 |

## Results

### Adequacy Test

For a total of 234 trees, all six initial models were successfully run. Among these, according to the corrected p-values (Table 1), all six models had quite comparable numbers of trees for which they were adequate (82.5-89.3%), with the exception of the diversity dependent model with both λ and μ depending linearly on K being adequate for fewer trees (73%). The 2D convex hulls showed a similar picture (adequate for 78.2-87.6%), with Yule, constant-rate birth-death, and λ-linear μ-constant diversity dependent doing slightly better than the other three models (Table 1).

Looking at the results from the perspective of trees, *all* models were adequate for a majority of them (61.1% or 58.1% respectively), with the percentage of trees for which fewer models were adequate declining quickly (Table 2). In cases where only one model was adequate, it was either the λ-exponential μ-constant time dependent (66.6%/42.9%), or the λ-linear μ-constant diversity dependent model (33.3%, more clearly so for 2D convex-hulls with 42.9%). When only one model was inadequate, it was usually the λ-linear μ-linear diversity dependent model according to the corrected p-values (62.9%), and the λ-exponential μ-constant time dependent model according to 2D convex-hulls (46.3%). For a small number of trees (12 according to corrected p-values, 5 according to 2D-convex-hulls), none of the six initial models were adequate (see below).

**Table 2:** Model Inadequacy by Tree. Number and percentage of trees for which a certain number of models were either adequate or inadequate.

| # Models | 0 | 1 | 2 | 3 | 4 | 5 | 6 |
| --- | --- | --- | --- | --- | --- | --- | --- |
| **Bonferroni-Pvalues** |  |  |  |  |  |  |  |
| **adequate** | 12 | 3 | 7 | 12 | 22 | 35 | 143 |
| **inadequate** | 143 | 35 | 22 | 12 | 7 | 3 | 12 |
| **% adequate** | 5.128 | 1.282 | 2.991 | 5.128 | 9.402 | 14.957 | 61.111 |
| **% inadequate** | 61.111 | 14.957 | 9.402 | 5.128 | 2.991 | 1.282 | 5.128 |
| **2D convex-hulls** |  |  |  |  |  |  |  |
| **adequate** | 5 | 7 | 15 | 14 | 16 | 41 | 136 |
| **inadequate** | 136 | 41 | 16 | 14 | 15 | 7 | 5 |
| **% adequate** | 2.137 | 2.991 | 6.410 | 5.983 | 6.838 | 17.521 | 58.120 |
| **% inadequate** | 58.120 | 17.521 | 6.838 | 5.983 | 6.410 | 2.991 | 2.137 |

When comparing how often combinations of models are inadequate (Table 3), both assessments agree that Yule and constant rate birth-death highly associated (in case of p-values, Yule is always inadequate when birth-death is), and both are often inadequate together with the λ-constant μ-exponential time dependent model, but less so vice-versa. Finally, when the λ-linear μ-constant diversity dependent model is inadequate, often the λ-linear μ-linear diversity dependent model is as well, but not vice-versa.

**Table 3:**
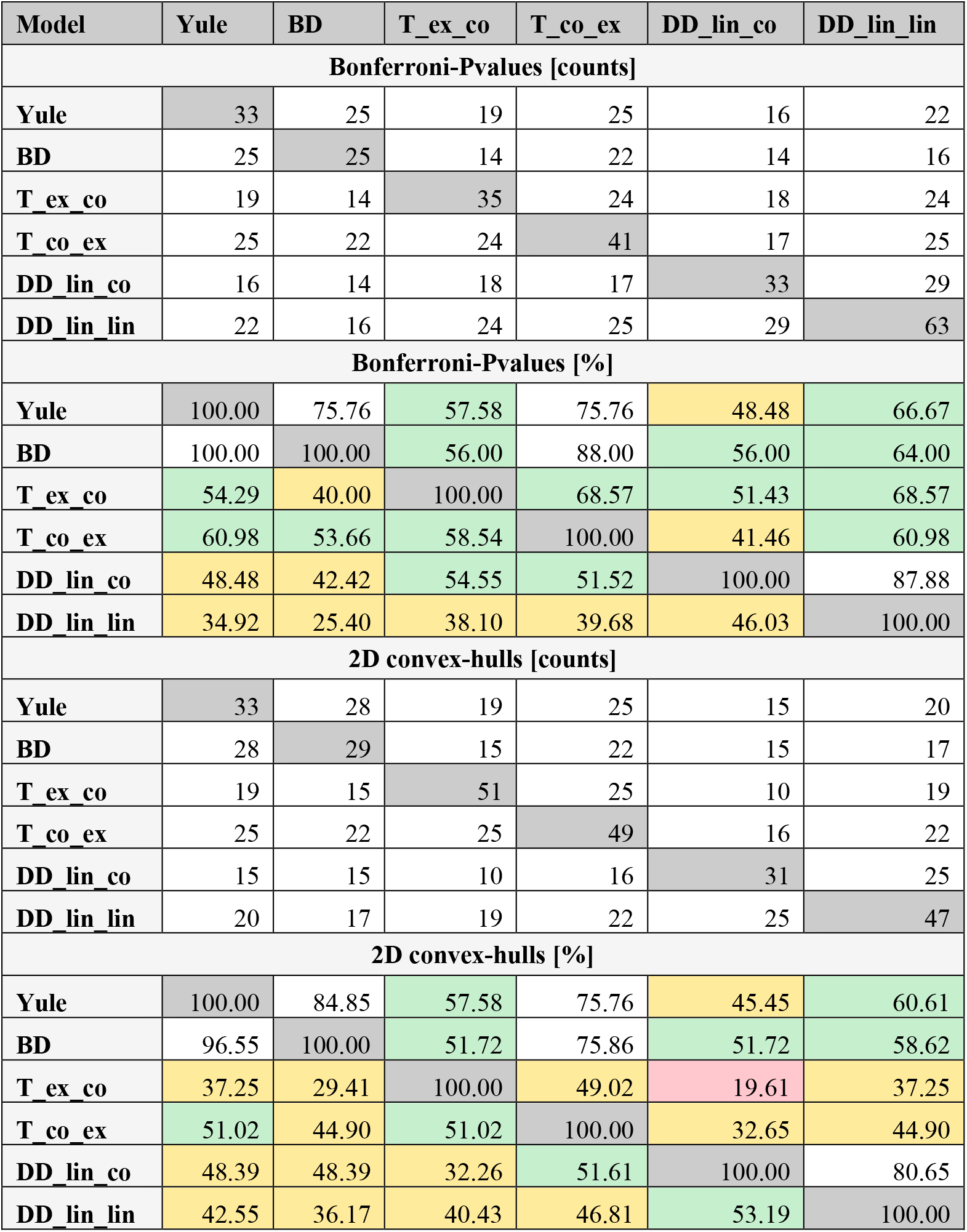
Pairwise Model Inadequacy. Pairwise counts of trees for which two models were inadequate for, and percentages of how often a model was inadequate together with another (by row), assessed using either Bonferroni-corrected p-values (above) or 2D convex-hulls (below). BD=constant rate birth-death, T_ex_co=time-dependent birth-death with exponential λ and constant μ, T_co_ex= time-dependent birth-death with constant λ and exponential μ. DD_lin_co= diversity-dependent birth-death with linear λ and constant μ. DD_lin_lin= diversity-dependent birth-death with linear λ and linear μ.

| Model | Yule | BD | T_ex_co | T_co_ex | DD_lin_co | DD_lin_lin |
| --- | --- | --- | --- | --- | --- | --- |
| **Bonferroni-Pvalues [counts]** |  |  |  |  |  |  |
| **Yule** | 33 | 25 | 19 | 25 | 16 | 22 |
| **BD** | 25 | 25 | 14 | 22 | 14 | 16 |
| **T_ex_co** | 19 | 14 | 35 | 24 | 18 | 24 |
| **T_co_ex** | 25 | 22 | 24 | 41 | 17 | 25 |
| **DD_lin_co** | 16 | 14 | 18 | 17 | 33 | 29 |
| **DD_lin_lin** | 22 | 16 | 24 | 25 | 29 | 63 |
| **Bonferroni-Pvalues [%]** |  |  |  |  |  |  |
| **Yule** | 100.00 | 75.76 | 57.58 | 75.76 | 48.48 | 66.67 |
| **BD** | 100.00 | 100.00 | 56.00 | 88.00 | 56.00 | 64.00 |
| **T_ex_co** | 54.29 | 40.00 | 100.00 | 68.57 | 51.43 | 68.57 |
| **T_co_ex** | 60.98 | 53.66 | 58.54 | 100.00 | 41.46 | 60.98 |
| **DD_lin_co** | 48.48 | 42.42 | 54.55 | 51.52 | 100.00 | 87.88 |
| **DD_lin_lin** | 34.92 | 25.40 | 38.10 | 39.68 | 46.03 | 100.00 |
| **2D convex-hulls [counts]** |  |  |  |  |  |  |
| **Yule** | 33 | 28 | 19 | 25 | 15 | 20 |
| **BD** | 28 | 29 | 15 | 22 | 15 | 17 |
| **T_ex_co** | 19 | 15 | 51 | 25 | 10 | 19 |
| **T_co_ex** | 25 | 22 | 25 | 49 | 16 | 22 |
| **DD_lin_co** | 15 | 15 | 10 | 16 | 31 | 25 |
| **DD_lin_lin** | 20 | 17 | 19 | 22 | 25 | 47 |
| **2D convex-hulls [%]** |  |  |  |  |  |  |
| **Yule** | 100.00 | 84.85 | 57.58 | 75.76 | 45.45 | 60.61 |
| **BD** | 96.55 | 100.00 | 51.72 | 75.86 | 51.72 | 58.62 |
| **T_ex_co** | 37.25 | 29.41 | 100.00 | 49.02 | 19.61 | 37.25 |
| **T_co_ex** | 51.02 | 44.90 | 51.02 | 100.00 | 32.65 | 44.90 |
| **DD_lin_co** | 48.39 | 48.39 | 32.26 | 51.61 | 100.00 | 80.65 |
| **DD_lin_lin** | 42.55 | 36.17 | 40.43 | 46.81 | 53.19 | 100.00 |

Inspection of the metrics shows that very commonly the skewness was responsible for the inadequacy of all models (Table 4). For Yule and constant-rate birth-death, it is followed by the principal Eigenvalue, while for exponential-λ constant-μ time-dependence and the two diversity dependent models, skewness was closely followed by peakedness. The constant-λ exponential-μ time-dependent model was mostly inadequate due to the principal Eigenvalue, followed by skewness. For the constant-λ exponential-μ time-dependent and the linear λ and μ diversity dependent model, the three metrics were more equally responsible than for the other four models.

**Table 4:** Metrics Discrepancy Between Adequate and Inadequate Models. Difference between number of taxa, tree shape metrics, and Eigengap between an empirical tree and the tree set simulated based on them under the respective models. The rows with the model names indicate for the three shape metrics, how often they were significantly different (and thus the reason the model was deemed inadequate) according to Bonferroni-corrected p-values.

| **Model (times inadequate)** | **Ntax** | **Princ. Eigenv.** | **Skewness** | **Peakedness** | **Eigengap** |
| --- | --- | --- | --- | --- | --- |
| **Yule (33)** |  | **11** | **22** | **3** |  |
| Mean Diff | 7.194 | 6504.625 | -0.522 | 0.510 | 32.087 |
| Mean Diff Ad. | -6.024 | -993.316 | -0.299 | 0.343 | 16.363 |
| Mean Diff Inad. | 87.704 | 52173.901 | -1.884 | 1.529 | 127.857 |
| **BD (25)** |  | **11** | **14** | **0** |  |
| Mean Diff | -6.834 | -1333.229 | -0.429 | 0.401 | 18.180 |
| Mean Diff Ad. | -8.781 | -1739.287 | -0.318 | 0.344 | 14.869 |
| Mean Diff Inad. | 9.444 | 2061.411 | -1.349 | 0.880 | 45.862 |
| **T_ex_co (35)** |  | **2** | **25** | **12** |  |
| Mean Diff | 41.717 | 9023.771 | -0.363 | 1.135 | 36.268 |
| Mean Diff Ad. | 35.693 | 7746.381 | -0.073 | 1.082 | 30.443 |
| Mean Diff Inad. | 75.967 | 16286.646 | -2.011 | 1.439 | 69.388 |
| **T_co_ex (41)** |  | **30** | **24** | **18** |  |
| Mean Diff | 10.576 | 6701.896 | -0.490 | 0.463 | 27.750 |
| Mean Diff Ad. | -0.768 | 3321.873 | -0.238 | 0.397 | 19.213 |
| Mean Diff Inad. | 63.976 | 22612.734 | -1.674 | 0.776 | 67.937 |
| **DD_lin_co (33)** |  | **2** | **24** | **15** |  |
| Mean Diff | 4.4276 | 1371.9054 | -0.8158 | 0.4076 | 25.6784 |
| Mean Diff Ad. | 2.2625 | 263.6969 | -0.6035 | 0.2952 | 20.4348 |
| Mean Diff Inad. | 17.6151 | 8121.9021 | -2.1085 | 1.0928 | 57.6166 |
| **DD_lin_lin (63)** |  | **18** | **46** | **31** |  |
| Mean Diff | 2.9490 | 313.8179 | -0.8562 | 0.2622 | 24.3850 |
| Mean Diff Ad. | 1.4881 | 74.2374 | -0.5744 | 0.2121 | 18.2381 |
| Mean Diff Inad. | 6.9143 | 964.1081 | -1.6212 | 0.3980 | 41.0696 |

In terms of actual metric values (Table 4), skewness tends to be lower in the simulated trees than in their empirical counterparts, meaning the simulated trees are tippier. This is true for all models, and whether or not they are adequate for a tree or not, with the difference being larger in trees modeled under inadequate models, and larger for trees for which more models are inadequate. Another global trend is that trees simulated under inadequate models tend to have a higher difference in principal Eigenvalue than those simulated under adequate models – in case of Yule and constant-rate birth-death, simulations under adequate models even have a lower principal Eigenvalue than the initial tree. This pattern is mirrored by the number of taxa in the trees. Also, peakedness and Eigengap (a measure of number of peaks in the spectrum, and indicator for how many distinct modes of diversification there are) while in general higher in all simulations, tend to be higher in trees simulated under an inadequate model.

We compared the metrics of the empirical trees between trees for which different numbers of models are adequate using an ANOVA and Tukey’s post-hoc test (Figure 1). Apart from slight patterns in age, principal Eigenvalue and Eigengap, the most striking signal comes from skewness, which seems to increase the chance that more models are inadequate for a tree if its skewness is more positive, meaning if the trees are very tippy.

**Figure 1:**
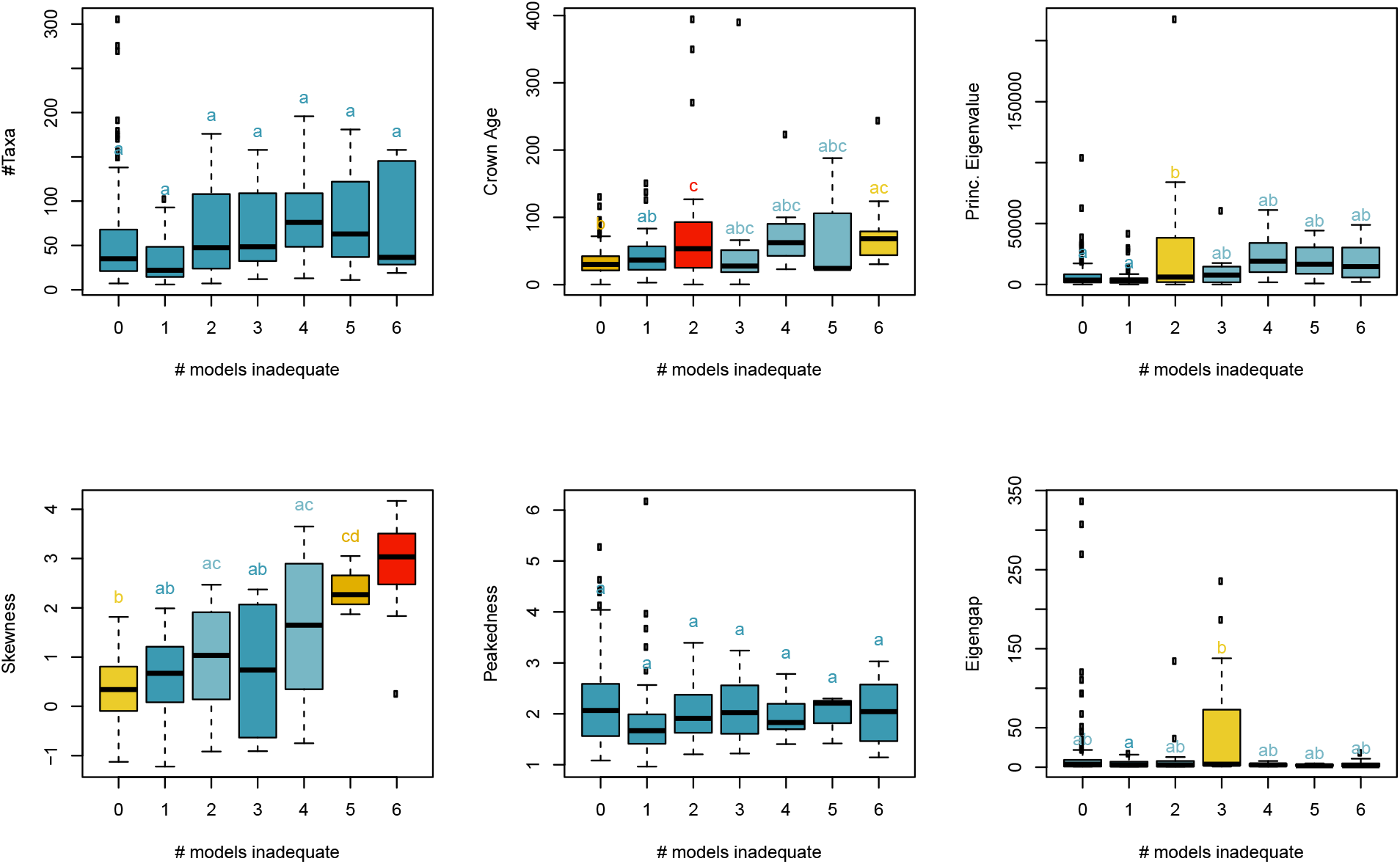
Relation of Empirical Tree Metrics with Model Inadequacy. Differences in number of taxa, crown age, and tree metrics for trees for which different numbers of the initial six models were inadequate. Colors and letters indicate the groupings based on a Tukey post-hoc test.

A comparison of tree metrics for which a particular model is adequate or inadequate (Table 5) shows that trees for which any model is inadequate tend to have a larger skewness than those for which the model is adequate. For Yule, birth-death, and linear-λ constant-μ density dependent, trees for which they are inadequate tend to additionally be larger, older, and have higher principal Eigenvalues – and a lower Eigengap in case of the diversity dependent model. Trees for which constant-λ exponential-μ time dependent models tend to be older than those for which it is adequate, as are trees for which linear-λ and μ is inadequate, with the addition that the latter also have a lower Eigengap. Finally, only for the exponential-λ constant-μ model there is a significant effect in peakedness (apart from the usual higher asymmetry and lower Eigengap), in that trees for which the model is inadequate have a lower peakedness.

**Table 5:** T-Tests of Metrics Between Adequate and Inadequate Models. Test for the difference between number of taxa, crown age, the tree shape metrics and Eigengap, between trees that were adequate and inadequate for a given model.

| Model | Attribute | Statistic | Parameter | Pvalue | ConfInt1 | ConfInt2 | Est1 | Est2 |
| --- | --- | --- | --- | --- | --- | --- | --- | --- |
| Yule | Ntax | -3.403 | 40.718 | **0.0015** | -54.972 | -14.021 | 50.413 | 84.909 |
|  | Tree Age | -3.174 | 33.776 | **0.0032** | -85.731 | -18.795 | 39.658 | 91.921 |
|  | Princ. Eigenv. | -2.948 | 33.180 | **0.0058** | -34523.310 | -6331.403 | 8181.396 | 28608.753 |
|  | Skewness | -3.936 | 34.146 | **0.0004** | -1.773 | -0.566 | 0.506 | 1.675 |
|  | Peakedness | -1.419 | 55.434 | 0.1616 | -0.411 | 0.070 | 2.104 | 2.275 |
|  | Eigengap | -1.097 | 37.066 | 0.2795 | -32.911 | 9.785 | 13.164 | 24.727 |
| BD | Ntax | -2.951 | 28.324 | **0.0063** | -59.777 | -10.809 | 51.507 | 86.800 |
|  | Tree Age | -2.477 | 25.594 | **0.0202** | -79.652 | -7.377 | 42.380 | 85.895 |
|  | Princ. Eigenv. | -2.977 | 29.068 | **0.0058** | -22644.131 | -4203.324 | 9628.018 | 23051.746 |
|  | Skewness | -2.277 | 25.243 | **0.0315** | -1.598 | -0.081 | 0.581 | 1.420 |
|  | Peakedness | -0.988 | 35.191 | 0.3298 | -0.416 | 0.143 | 2.114 | 2.250 |
|  | Eigengap | -1.375 | 26.066 | 0.1807 | -46.148 | 9.145 | 12.818 | 31.320 |
| T_ex_co | Ntax | 0.671 | 50.395 | 0.5053 | -11.499 | 23.038 | 56.141 | 50.371 |
|  | Tree Age | -1.896 | 38.623 | 0.0654 | -52.043 | 1.685 | 43.263 | 68.442 |
|  | Princ. Eigenv. | -1.038 | 52.750 | 0.3042 | -9998.894 | 3181.706 | 10552.345 | 13960.939 |
|  | Skewness | -7.373 | 38.529 | **0.0000** | -2.042 | -1.162 | 0.431 | 2.033 |
|  | Peakedness | 2.898 | 58.147 | **0.0053** | 0.108 | 0.590 | 2.181 | 1.831 |
|  | Eigengap | 3.852 | 211.673 | **0.0002** | 6.295 | 19.495 | 16.724 | 3.829 |
| T_co_ex | Ntax | -0.525 | 54.380 | 0.6014 | -23.867 | 13.954 | 54.409 | 59.366 |
|  | Tree Age | -2.481 | 47.035 | **0.0167** | -53.118 | -5.551 | 41.889 | 71.224 |
|  | Princ. Eigenv. | -1.476 | 68.935 | 0.1444 | -10539.535 | 1574.252 | 10276.757 | 14759.399 |
|  | Skewness | -5.569 | 44.967 | **0.0000** | -1.720 | -0.806 | 0.449 | 1.713 |
|  | Peakedness | 1.740 | 75.407 | 0.0859 | -0.029 | 0.428 | 2.163 | 1.964 |
|  | Eigengap | -0.321 | 52.513 | 0.7493 | -19.369 | 14.022 | 14.326 | 17.000 |
| DD_lin_co | Ntax | -2.168 | 42.687 | **0.0358** | -40.302 | -1.456 | 52.333 | 73.212 |
|  | Tree Age | -2.486 | 35.172 | **0.0178** | -65.819 | -6.645 | 41.919 | 78.151 |
|  | Princ. Eigenv. | -2.960 | 43.189 | **0.0050** | -18689.774 | -3542.383 | 9494.525 | 20610.604 |
|  | Skewness | -5.719 | 37.200 | **0.0000** | -1.734 | -0.827 | 0.490 | 1.770 |
|  | Peakedness | 1.804 | 62.352 | 0.0760 | -0.022 | 0.422 | 2.157 | 1.957 |
|  | Eigengap | 2.978 | 231.793 | **0.0032** | 3.512 | 17.248 | 16.259 | 5.879 |
| DD_lin_lin | Ntax | -0.431 | 112.771 | 0.6675 | -17.943 | 11.536 | 54.415 | 57.619 |
|  | Tree Age | -2.103 | 85.958 | **0.0384** | -36.712 | -1.033 | 41.948 | 60.820 |
|  | Princ. Eigenv. | -1.611 | 73.612 | 0.1114 | -14285.236 | 1510.953 | 9342.562 | 15729.704 |
|  | Skewness | -5.581 | 77.529 | **0.0000** | -1.314 | -0.623 | 0.410 | 1.378 |
|  | Peakedness | 1.293 | 118.487 | 0.1984 | -0.080 | 0.380 | 2.169 | 2.019 |
|  | Eigengap | 3.692 | 181.837 | **0.0003** | 6.644 | 21.901 | 18.637 | 4.365 |

### Trees without adequate Models

Among trees for which all models ran, no model was adequate for twelve trees according to the Bonferroni-corrected p-values, and for five trees according to the 2D convex-hulls, with an overlap of four trees for which no model was adequate according to either way of assessing it. When running the additional set of models on those four trees, adequate models were found for two of them, while the other two still could not be adequately described by any model used (however, about half of the models used on those trees did not run successfully).

In terms of tree metrics, the two trees for which all models are inadequate, do not stand out in any particular way (Figure 2), apart from their high skewness, as expected based on the results above. However, this would not explain what differentiates these two trees from the ones for which only the six initial models were inadequate, but for which an adequate one was found among the additional eleven models. The inferred phylogenies and associated Laplacian spectra (Figure 3) suggest that the source of the high skewness might be that these trees have a small secondary peak at higher Eigenvalues than the main peak. These seem to result from the species/clade that is sister to the rest of each respective tree, and that they are not only subtending a for this tree comparatively long branch, but that the rest of the tree is subtending a relatively long branch as well.

**Figure 2:**
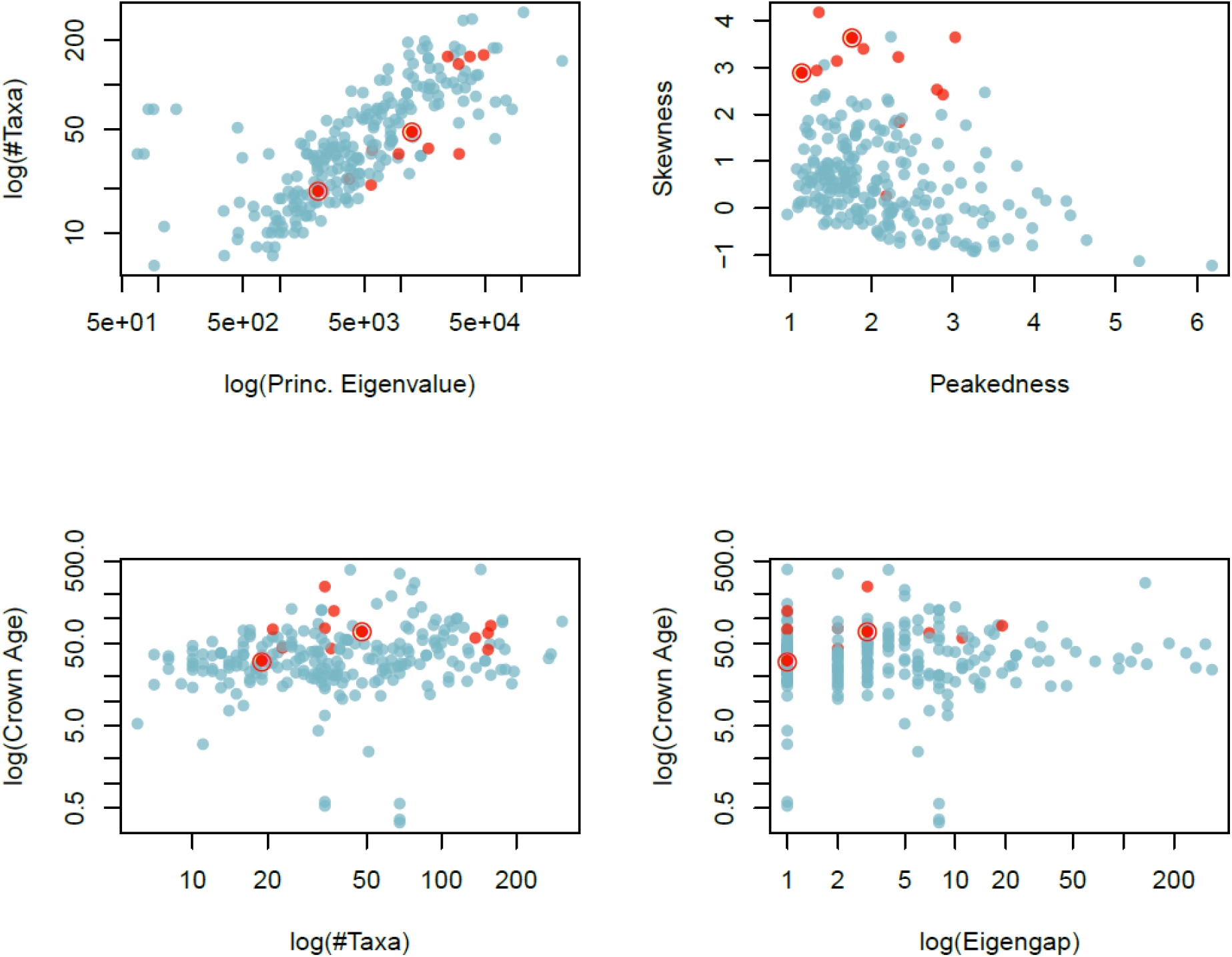
Empirical Tree Metrics and Inadequate Models. Scatterplots of number of taxa, crown ages and tree metrics, color coded by the number of models inadequate for each tree. Blue indicates at least one of the initial six models was adequate, red indicates all six were inadequate, and the two trees for which every model was inadequate are marked with red circles. The axes for number of taxa, crown age, principal Eigenvalue, and Eigengap are logarithmized.

**Figure 3:**
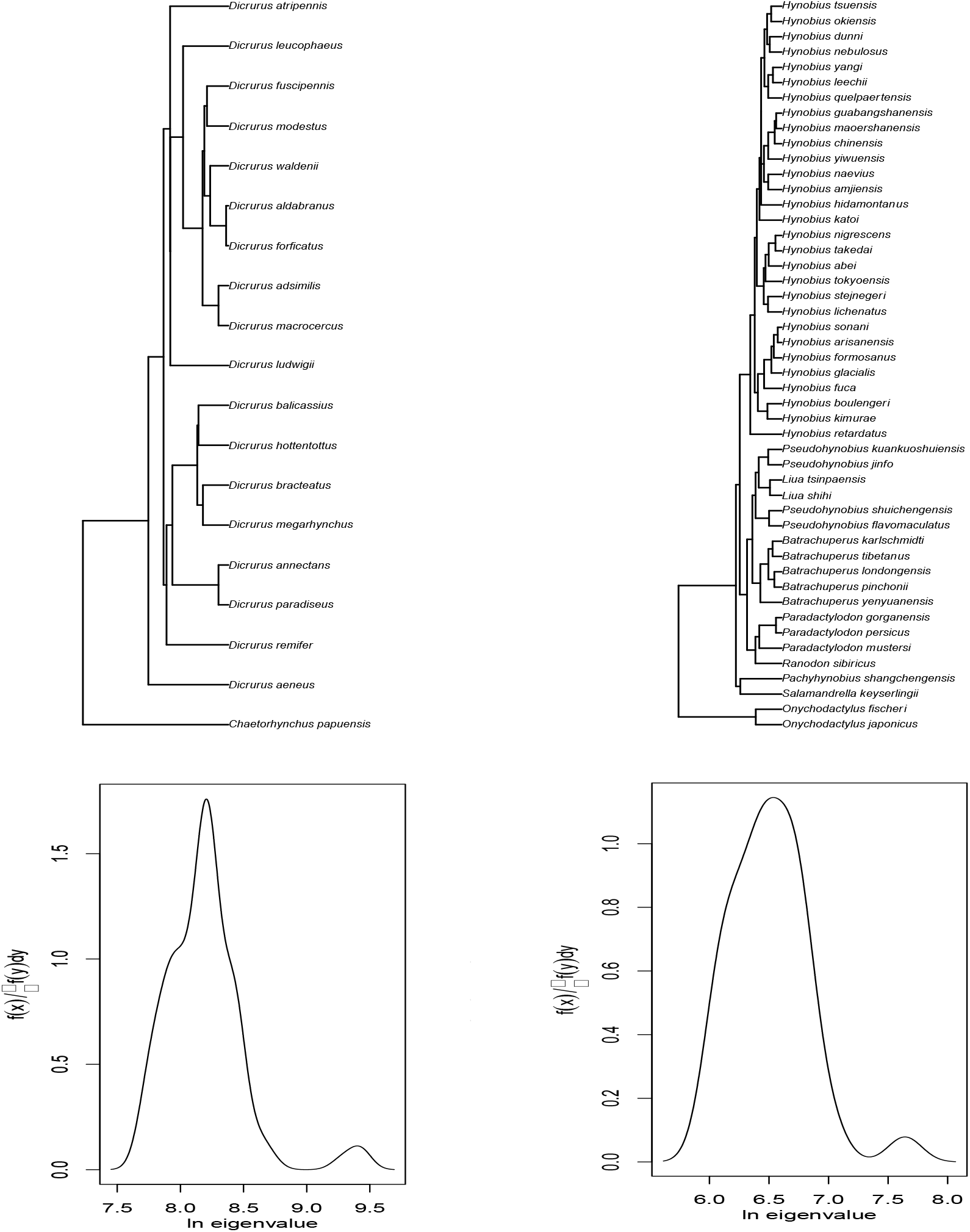
Phylogenies and Spectra of Trees for Which All Models Fail. The phylogenies (top) and corresponding Laplacian spectra (bottom) of the two trees for which all models are inadequate. On the left tree 244 (family Dricuridae), on the right tree 323 (family Hynobiidae).

### Adequacy and Fit

Overall, of 234 trees for which models were fitted, in 198 cases the best fitting model was adequate. In 22 cases the best model was inadequate, but at least one of the other models was adequate, and for 14 trees none of those four models were adequate (Figure 4).

**Figure 4:**
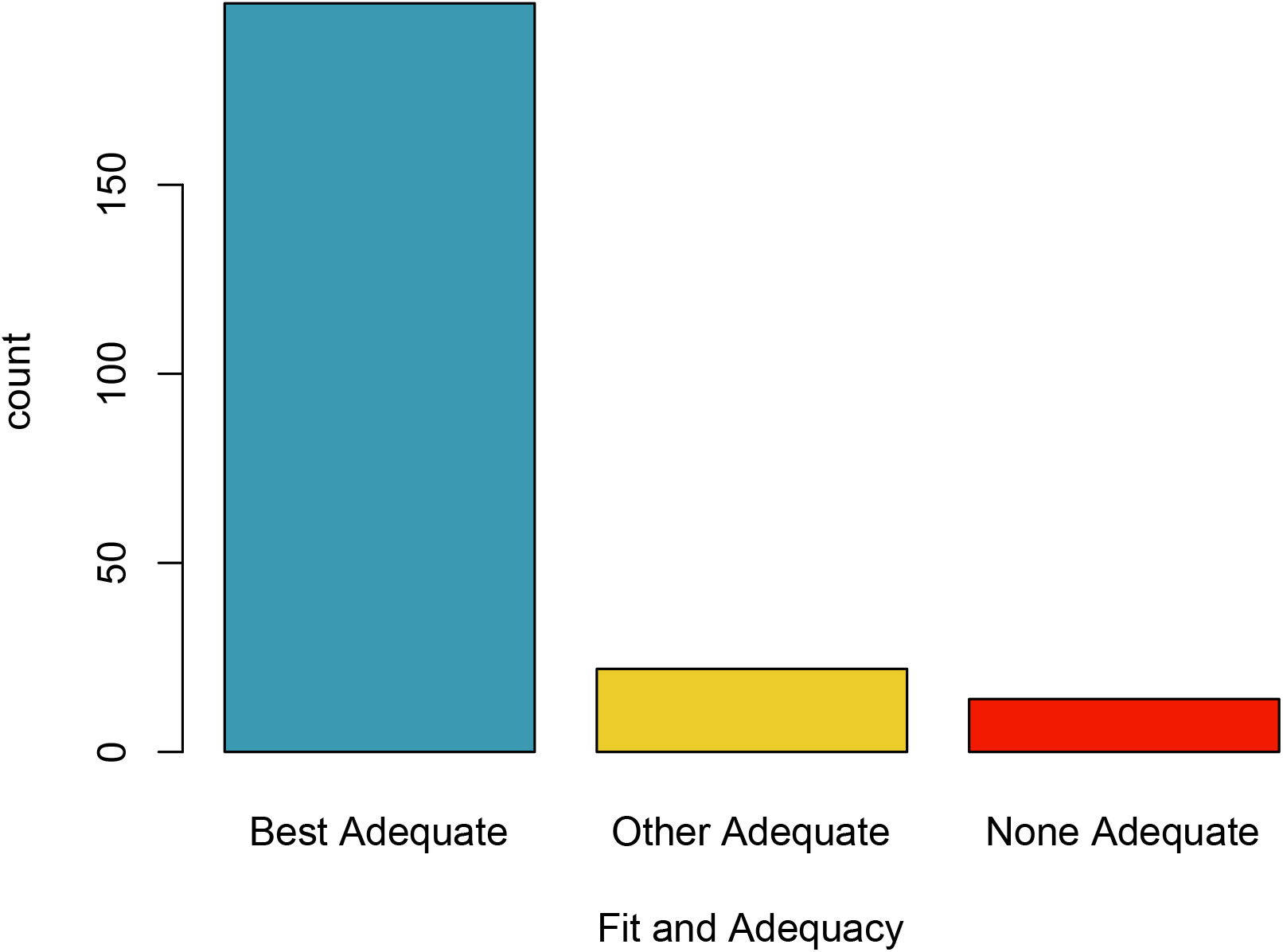
Relation of Model Fit and Model Adequacy. Numbers of trees for which their best fitting and other model was adequate, a different model than the best was adequate, or none were adequate.

## Discussion

We have seen a steady increase of models used to study diversification, and the equally increasing awareness of the issues and shortcomings those models have. In the light of this, steps need to be taken towards gaining insight into whether and how these issues affect the results inferred under these models, thereby both improving our inference and increasing confidence in our findings. This research addresses this issue by applying a new method to test the adequacy of diversification models to a large set of empirical phylogenies, in order to uncover patterns in model adequacy and the causes behind them.

### Overall Adequacy Patterns

Overall, the results draw a promising picture of the adequacy of our models (assuming the adequacy test is accurate, and the metrics used as test statistics capture meaningful aspects of the trees). All models under investigation were adequate for a relatively high number of trees, and also from the tree’s perspective, not only were all or many models adequate for most trees, even with the modest set of six basic models, for only a small number of trees could an adequate model not be found. It might come at a surprise that the models performed so evenly across the trees, as one could have imagined that *e.g.* overly simplistic models like Yule would only be adequate for a particular kind of tree. However, this seems not to be the case. This presumably demonstrates the flexibility of those models, partially resulting from the stochasticity underlying the processes – even when we use a model with one constant rate of lineage accumulation, a relatively wide range of branching patterns can still emerge from it by chance (Slowinski & Guyer 1989). However, it calls into question whether such a model has too much stochasticity to allow reliable inference, or in our case of adequacy testing. The wide range of tree shape space a model covers may make it seem adequate without necessarily meaning that the model really tells us anything reliable about the diversification process underlying a certain tree.

Models that are adequate when the others fail (λ-exponential μ-constant time dependent or λ-linear μ-constant diversity dependent), might suggest that these models are able to adequately describe particular branching patterns, for which the stochasticity of the other models could not account. For example, exponential variation in λ should be able to account for more extreme cases of increases or decreases in diversification on either extreme of time, whereas the diversity dependence model would allow for the kind of stagnation other models could not account for. Indeed, the challenge of being able to detect decreases in diversification rates have been noted previously (Liow, Quental & Marshall 2010; Burin *et al.* 2019).

### Trees without adequate Models

It is both concerning and exciting to find phylogenies for which no model is adequate. The concern arises because the field lacks the tools to address come case studies. But those cases might reveal new aspects of diversification and drive the exploration of new models. Particularly with the limited number of models used here – and demonstrated by applying the extended set of models to the initially unmatched trees – it is conceivable that the adequate model for those cases might indeed already exist and is just waiting to be employed.

Alternatively, it is possible that the tree shape metrics used here are not appropriate – or not all of the appropriate necessary metrics – to use. Inadequate models maybe deemed adequate if these metrics fail to capture a crucial aspect of the trees lost on the model. Adequate models may seem inadequate if the metrics capture variation in an aspect of the tree that is not actually related to its underlying diversification process. However, while additional and alternative metrics should still be explored in the future, there is at least some confidence in the ones used here, as they have generally been shown to be able to distinguish different tree shapes (Lewitus & Morlon 2016a).

Another possibility, given that the trees in question only represent less than 1% of the tested trees, is that *BoskR* erroneously deemed models inadequate for them. This would relegate these un-matchable trees simply be the product of type-I error. As mentioned before, there is a certain amount of stochasticity involved in the process, and as discussed in Schwery and O’Meara (2020), the results can be significantly influenced by things like *e.g.* insufficient simulations to generate a distribution of shape metrics that properly represents the properties of a certain pairing of model and parameters.

Returning to the two actual trees for which no model was adequate, their high value for skewedness appears to be the only aspect that, at least to some extent, differentiates them from trees for which models were found adequate. As it appears that the comparatively large amount of branch length that separates the bulk of the species from those sister to them (Figure 3), the question arises whether these trees might simply include their outgroup. Tree 244 is of the bird family Dricuridae, tree 323 of the salamander family Hynobiidae. While the suspected outgroup of the Hynobiidae (*Onychodactylus*) is indeed considered a part of the family, that of the Dricuridae, *Chaetorhynchus papuensis,* has recently been reclassified as belonging to the family Rhipiduridae (Barker *et al.* 2004; Irestedt *et al.* 2008), meaning it might represent something of an unintended outgroup, which explains its distance to the rest of the group.

Finally, there is an implicit assumption in this approach, the violation of which could have a large impact on the validity of its results. We are assuming that the empirical trees are correct representations of the true trees (or a close enough representation thereof). Any model could, in theory, perfectly describe the diversification dynamics of a group of organisms but would still be marked inadequate if the empirical tree we judge it by does not represent the true diversification patterns of that group. On the more specific and technical side of this argument (and of the term ‘correct’), we are assuming that there is no relevant bias on the tree shape stemming from the process that we employed to infer the tree. Even if no mistakes in the strict sense are made, biases in tree inference or divergence time estimation could have a large effect on the shape of the tree, and it seems possible that *e.g.* the use of a Yule or birth-death prior in BEAST could bias a tree towards looking like those models would be adequate for it. And lastly, while these models account for past diversity in now extinct lineages (that are thus missing from the tree) by estimating extinction rates, it is known that getting accurate estimates is possible, but challenging (Rabosky 2010; Beaulieu & O’Meara 2015; Rabosky 2016). Thus, without knowledge and explicit inclusion of past diversity in the trees, a degree of uncertainty of the models’ actual adequacy will remain.

### Relation of Adequacy and Model Fit

The finding that model fit and model adequacy do not necessarily coincide – or in other words, that an adequate model does not necessarily fit the data better than an inadequate one – is important in two ways. It demonstrates that the practice of model selection based on best fit is insufficient and might lead to false results. Adding adequacy testing to the procedure facilitates the consideration of only those models that provide meaningful results. Additionally, our example suggests the utility of adequacy testing in another way: there are cases when all models implemented in a certain framework (*e.g.* a specific R package) are inadequate. Knowing this would encourage a researcher to venture out and explore the models implemented elsewhere.

## Acknowledgements

We would like to thank Luna Sanchez-Reyes, Jeremy Beaulieu, Dan Simberloff, and Colin Sumrall for helpful discussions and comments at various stages of this project.

## References

Alfaro, M.E., Santini, F., Brock, C., Alamillo, H., Dornburg, A., Rabosky, D.L., Carnevale, G. & Harmon, L.J. (2009) Nine exceptional radiations plus high turnover explain species diversity in jawed vertebrates. Proceedings of the National Academy of Sciences of the United States of America, 106, 13410–13414.

Barker, F.K., Cibois, A., Schikler, P., Feinstein, J. & Cracraft, J. (2004) Phylogeny and diversification of the largest avian radiation. Proceedings of the National Academy of Sciences, 101, 11040–11045.

Beaulieu, J.M. & O’Meara, B.C. (2015) Extinction can be estimated from moderately sized molecular phylogenies. Evolution, 69, 1036–1043.

Beaulieu, J.M. & O’Meara, B.C. (2016) Detecting Hidden Diversification Shifts in Models of Trait-Dependent Speciation and Extinction. Systematic Biology, 65, 583–601.

Bouchenak-Khelladi, Y., Muasya, A.M. & Linder, H.P. (2014) A revised evolutionary history of Poales: origins and diversification. Botanical Journal of the Linnean Society, 175, 4–16.

Brown, J.M. & ElDabaje, R. (2008) PuMA: Bayesian analysis of p artitioned (and u npartitioned) m odel a dequacy. Bioinformatics, 25, 537–538.

Burin, G., Alencar, L.R.V., Chang, J., Alfaro, M.E. & Quental, T.B. (2019) How Well Can We Estimate Diversity Dynamics for Clades in Diversity Decline? Systematic Biology, 68, 47–62.

Caetano, D.S., O’Meara, B.C. & Beaulieu, J.M. (2018) Hidden state models improve state-dependent diversification approaches, including biogeographical models. Evolution, 72, 2308–2324.

FitzJohn, R.G. (2010) Quantitative Traits and Diversification. Systematic Biology, 59, 619–633.

FitzJohn, R.G. (2012) Diversitree: comparative phylogenetic analyses of diversification in R. Methods in Ecology and Evolution, 3, 1084–1092.

FitzJohn, R.G., Maddison, W.P. & Otto, S.P. (2009) Estimating trait-dependent speciation and extinction rates from incompletely resolved phylogenies. Systematic Biology, 58, 595–611.

Goldberg, E.E., Lancaster, L.T. & Ree, R.H. (2011) Phylogenetic Inference of Reciprocal Effects between Geographic Range Evolution and Diversification. Systematic Biology, 60, 451–465.

Hinchliff, C.E., Smith, S.A., Allman, J.F., Burleigh, J.G., Chaudhary, R., Coghill, L.M., Crandall, K.A., Deng, J., Drew, B.T., Gazis, R., Gude, K., Hibbett, D.S., Katz, L.A., Laughinghouse, H.D., McTavish, E.J., Midford, P.E., Owen, C.L., Ree, R.H., Rees, J.A., Soltis, D.E., Williams, T. & Cranston, K.A. (2015) Synthesis of phylogeny and taxonomy into a comprehensive tree of life. Proceedings of the National Academy of Sciences of the United States of America, 112, 12764–12769.

Höhna, S., May, M.R. & Moore, B.R. (2015) TESS: an R package for efficiently simulating phylogenetic trees and performing Bayesian inference of lineage diversification rates. Bioinformatics, 32, 789–791.

Irestedt, M., Fuchs, J., Jonsson, K.A., Ohlson, J.I., Pasquet, E. & Ericson, P.G. (2008) The systematic affinity of the enigmatic Lamprolia victoriae (Aves: Passeriformes)--an example of avian dispersal between New Guinea and Fiji over Miocene intermittent land bridges? Molecular Phylogenetics and Evolution, 48, 1218–1222.

Laenen, B., Shaw, B., Schneider, H., Goffinet, B., Paradis, E., Desamore, A., Heinrichs, J., Villarreal, J.C., Gradstein, S.R., McDaniel, S.F., Long, D.G., Forrest, L.L., Hollingsworth, M.L., Crandall-Stotler, B., Davis, E.C., Engel, J., Von Konrat, M., Cooper, E.D., Patino, J., Cox, C.J., Vanderpoorten, A. & Shaw, A.J. (2014) Extant diversity of bryophytes emerged from successive post-Mesozoic diversification bursts. Nature Communications, 5.

Lewitus, E. & Morlon, H. (2016a) Characterizing and Comparing Phylogenies from their Laplacian Spectrum. Systematic Biology, 65, 495–507.

Lewitus, E. & Morlon, H. (2016b) Natural Constraints to Species Diversification. Plos Biology, 14.

Liow, L.H., Quental, T.B. & Marshall, C.R. (2010) When can decreasing diversification rates be detected with molecular phylogenies and the fossil record? Systematic Biology, 59, 646–659.

Maddison, W.P. & FitzJohn, R.G. (2015) The Unsolved Challenge to Phylogenetic Correlation Tests for Categorical Characters. Systematic Biology, 64, 127–136.

Maddison, W.P., Midford, P.E. & Otto, S.P. (2007) Estimating a binary character’s effect on speciation and extinction. Systematic Biology, 56, 701–710.

Nguyen, V.D., Nguyen, T.H., Tayeen, A.S.M., Laughinghouse, H.D., Sánchez-Reyes, L.L., Pontelli, E., Mozzherin, D., O’Meara, B. & Stoltzfus, A. (2018) Phylotastic: improving access to tree-of-life knowledge with flexible, on-the-fly delivery of trees. BioRxiv, 419143.

O’Meara, B.C., Smith, S.D., Armbruster, W.S., Harder, L.D., Hardy, C.R., Hileman, L.C., Hufford, L., Litt, A., Magallón, S. & Smith, S.A. (2016) Non-equilibrium dynamics and floral trait interactions shape extant angiosperm diversity. Proceedings of the Royal Society B: Biological Sciences, 283, 20152304.

Pennell, M.W., FitzJohn, R.G., Cornwell, W.K. & Harmon, L.J. (2015) Model Adequacy and the Macroevolution of Angiosperm Functional Traits. American Naturalist, 186, E33–E50.

Pigot, A.L., Owens, I.P.F. & Orme, C.D.L. (2012) Speciation and Extinction Drive the Appearance of Directional Range Size Evolution in Phylogenies and the Fossil Record. Plos Biology, 10.

Rabosky, D.L. (2010) Extinction rates should not be estimated from molecular phylogenies. Evolution: International Journal of Organic Evolution, 64, 1816–1824.

Rabosky, D.L. (2014) Automatic detection of key innovations, rate shifts, and diversity-dependence on phylogenetic trees. Plos One, 9, e89543.

Rabosky, D.L. (2016) Challenges in the estimation of extinction from molecular phylogenies: A response to Beaulieu and O’Meara. Evolution, 70, 218–228.

Rabosky, D.L. & Goldberg, E.E. (2015) Model Inadequacy and Mistaken Inferences of Trait-Dependent Speciation. Systematic Biology, 64, 340–355.

Revell, L.J., Harmon, L.J. & Glor, R.E. (2005) Under-parameterized model of sequence evolution leads to bias in the estimation of diversification rates from molecular phylogenies. Systematic Biology, 54, 973–983.

Schwery O. & O’Meara B.C. (2020) BoskR-Testing Adequacy of Diversification Models Using Tree Shape. bioRxiv.:2020.12.21.423829.

Schwery, O., Onstein, R.E., Bouchenak-Khelladi, Y., Xing, Y., Carter, R.J. & Linder, H.P. (2015) As old as the mountains: the radiations of the Ericaceae. New Phytologist, 207, 355–367.

Slowinski, J.B. & Guyer, C. (1989) Testing the stochasticity of patterns of organismal diversity: an improved null model. The American Naturalist, 134, 907–921.

Stadler, T. (2013) How Can We Improve Accuracy of Macroevolutionary Rate Estimates? Systematic Biology, 62, 321–329.

Steeman, M.E., Hebsgaard, M.B., Fordyce, R.E., Ho, S.Y.W., Rabosky, D.L., Nielsen, R., Rahbek, C., Glenner, H., Sorensen, M.V. & Willerslev, E. (2009) Radiation of Extant Cetaceans Driven by Restructuring of the Oceans. Systematic Biology, 58, 573–585.

Stoltzfus, A., Lapp, H., Matasci, N., Deus, H., Sidlauskas, B., Zmasek, C.M., Vaidya, G., Pontelli, E., Cranston, K., Vos, R., Webb, C.O., Harmon, L.J., Pirrung, M., O’Meara, B., Pennell, M.W., Mirarab, S., Rosenberg, M.S., Balhoff, J.P., Bik, H.M., Heath, T.A., Midford, P.E., Brown, J.W., McTavish, E.J., Sukumaran, J., Westneat, M., Alfaro, M.E., Steele, A. & Jordan, G. (2013) Phylotastic! Making tree-of-life knowledge accessible, reusable and convenient. Bmc Bioinformatics, 14.

Xing, Y.W., Onstein, R.E., Carter, R.J., Stadler, T. & Linder, H.P. (2014) Fossils and a large molecular phylogeny show that the evolution of species richness, generic diversity, and turnover rates are disconnected. Evolution, 68, 2821–2832.

