## Supplementary Materials for "The Shape of Trees – Limits of Current Diversification Models"

**Supplementary Material for:**
**Schwery, O & O'Meara, BC (2021) The Shape of Trees – Limits of Current Diversification Models**

**Table S1: Collected Empirical Phylogenies.**
Tree ID given in this study, number of taxa represented, crown age, and citation of the original study that published each tree.

| TreeID | Taxa | Age | Source |
|---|---|---|---|
| Emp_1 | 87 | 35.86 | Steeman, M.E., Hebsgaard, M.B., Fordyce, R.E., Ho, S.Y.W., Rabosky, D.L., Nielsen, R., Rahbek, C., Glenner, H., Sorensen, M.V. & Willerslev, E. (2009) Radiation of Extant Cetaceans Driven by Restructuring of the Oceans. Systematic Biology, 58, 573-585. |
| Emp_2 | 77 | 117.28 | Schwery, O., Onstein, R.E., Bouchenak-Khelladi, Y., Xing, Y., Carter, R.J. & Linder, H.P. (2015) As old as the mountains: the radiations of the Ericaceae. New Phytologist, 207, 355-367 |
| Emp_3 | 478 | 122.19 | Schwery, O., Onstein, R.E., Bouchenak-Khelladi, Y., Xing, Y., Carter, R.J. & Linder, H.P. (2015) As old as the mountains: the radiations of the Ericaceae. New Phytologist, 207, 355-367 |
| Emp_4 | 557 | 124.13 | Bouchenak-Khelladi, Y., Muasya, A.M. & Linder, H.P. (2014) A revised evolutionary history of Poales: origins and diversification. Botanical Journal of the Linnean Society, 175, 4-16. |
| Emp_5 | 584 | 118.20 | Xing, Y.W., Onstein, R.E., Carter, R.J., Stadler, T. & Linder, H.P. (2014) Fossils and a large molecular phylogeny show that the evolution of species richness, generic diversity, and turnover rates are disconnected. Evolution, 68, 2821-2832 |
| Emp_6 | 549 | 136.00 | O'Meara BC, Smith SD, Armbruster WS, Harder LD, Hardy CR, Hileman LC, Hufford L, Litt A, Magallón S, Smith SA, Stevens PF. Non-equilibrium dynamics and floral trait interactions shape extant angiosperm diversity. Proceedings of the Royal Society B: Biological Sciences. 2016 May 11;283(1830):20152304. |
| Emp_7 | 309 | 536.16 | Laenen, B., Shaw, B., Schneider, H., Goffinet, B., Paradis, E., Desamore, A., Heinrichs, J., Villarreal, J.C., Gradstein, S.R., McDaniel, S.F., Long, D.G., Forrest, L.L., Hollingsworth, M.L., Crandall-Stotler, B., Davis, E.C., Engel, J., Von Konrat, M., Cooper, E.D., Patino, J., Cox, C.J., Vanderpoorten, A. & Shaw, A.J. (2014) Extant diversity of bryophytes emerged from successive post-Mesozoic diversification bursts. Nature Communications, 5 |
| Emp_8 | 9 | 16.17 | Steeman, M.E., Hebsgaard, M.B., Fordyce, R.E., Ho, S.Y.W., Rabosky, D.L., Nielsen, R., Rahbek, C., Glenner, H., Sorensen, M.V. & Willerslev, E. (2009) Radiation of Extant Cetaceans Driven by Restructuring of the Oceans. Systematic Biology, 58, 573-585. |
| Emp_9 | 11 | 2.99 | Pigot, A.L., Owens, I.P.F. & Orme, C.D.L. (2012) Speciation and Extinction Drive the Appearance of Directional Range Size Evolution in Phylogenies and the Fossil Record. Plos Biology, 10 |
| Emp_10 | 6 | 5.31 | Steeman, M.E., Hebsgaard, M.B., Fordyce, R.E., Ho, S.Y.W., Rabosky, D.L., Nielsen, R., Rahbek, C., Glenner, H., Sorensen, M.V. & Willerslev, E. (2009) Radiation of Extant Cetaceans Driven by Restructuring of the Oceans. Systematic Biology, 58, 573-585. |
| Emp_11 | 150 | 30.69 | Steeman, M.E., Hebsgaard, M.B., Fordyce, R.E., Ho, S.Y.W., Rabosky, D.L., Nielsen, R., Rahbek, C., Glenner, H., Sorensen, M.V. & Willerslev, E. (2009) Radiation of Extant Cetaceans Driven by Restructuring of the Oceans. Systematic Biology, 58, 573-585. |

Table S1 Continued

| TreeID | Taxa | Age | Source |
|---|---|---|---|
| Emp_12 | 52 | 50.50 | Selvatti, Alexandre Pedro, Ana Galvao, Anieli Guirro Pereira, Luiz Pedreira Gonzaga, Claudia Augusta de Moraes Russo. 2016. An African origin of the Eurylaimides (Passeriformes) and the successful diversification of the ground-foraging pittas (Pittidae). Molecular Biology and Evolution, p. msw250 |
| Emp_13 | 118 | 78.49 | Fine, Paul V. A., Felipe Zapata, Douglas C. Daly. 2014. Investigating processes of neotropical rain forest tree diversification by examining the evolution and historical biogeography of the Protieae (Burseraceae). Evolution 68 (7): 1988-2004 |
| Emp_14 | 76 | 28.48 | Arbabi, Tayebeh, Javier Gonzalez, Michael Wink. 2014. A re-evaluation of phylogenetic relationships within reed warblers (Aves: Acrocephalidae) based on eight molecular loci and ISSR profiles. Molecular Phylogenetics and Evolution 78: 304-313. |
| Emp_15 | 174 | 3.87 | Ralph S. Peters, Lars Krogmann, Christoph Mayer, Alexander Donath, Simon Gunkel, Karen Meusemann, Alexey Kozlov, Lars Podsiadlowski, Malte Petersen, Robert Lanfear, Patricia A. Diez, John Heraty, Karl M. Kjer, Seraina Klopfstein, Rudolf Meier, Carlo Polidori, Thomas Schmitt, Shanlin Liu, Xin Zhou, Torsten Wappler, Jes Rust, Bernhard Misof, Oliver Niehuis, 2017, 'Evolutionary History of the Hymenoptera', Current Biology |
| Emp_16 | 148 | 425.00 | A. E. Syme, T. H. Oakley, 2011, 'Dispersal between Shallow and Abyssal Seas and Evolutionary Loss and Regain of Compound Eyes in Cylindroleberidid Ostracods: Conflicting Conclusions from Different Comparative Methods', Systematic Biology, vol. 61, no. 2, pp. 314-336 |
| Emp_17 | 103 | 72.81 | Daniela Campanella, Lily C. Hughes, Peter J. Unmack, Devin D. Bloom, Kyle R. Piller, Guillermo Orti, 2015, 'Multi-locus fossil-calibrated phylogeny of Atheriniformes (Teleostei, Ovalentaria)', Molecular Phylogenetics and Evolution, vol. 86, pp. 8-23 |
| Emp_18 | 68 | 49.71 | Upham, N.S. & B.D. Patterson. 2015. Evolution of the caviomorph rodents: a complete phylogeny and timetree of living genera. Pp. 63-120 In: Biology of caviomorph rodents: diversity and evolution (A. I. Vassallo & D. Antenucci, eds.). SAREM Series A, Buenos Aires. |
| Emp_19 | 78 | 271.08 | Vea, Isabelle M., David A. Grimaldi. 2016. Putting scales into evolutionary time: the divergence of major scale insect lineages (Hemiptera) predates the radiation of modern angiosperm hosts. Scientific Reports, 6: 23487 . |
| Emp_20 | 88 | 15.20 | Hugall, Andrew F., Devi Stuart-Fox. 2012. Accelerated speciation in colour-polymorphic birds. Nature 485 (7400): 631-634. |
| Emp_21 | 111 | 30.22 | Hugall, Andrew F., Devi Stuart-Fox. 2012. Accelerated speciation in colour-polymorphic birds. Nature 485 (7400): 631-634. |
| Emp_22 | 198 | 39.87 | Hugall, Andrew F., Devi Stuart-Fox. 2012. Accelerated speciation in colour-polymorphic birds. Nature 485 (7400): 631-634. |
| Emp_23 | 53 | 27.86 | Hugall, Andrew F., Devi Stuart-Fox. 2012. Accelerated speciation in colour-polymorphic birds. Nature 485 (7400): 631-634. |
| Emp_24 | 181 | 24.22 | Hugall, Andrew F., Devi Stuart-Fox. 2012. Accelerated speciation in colour-polymorphic birds. Nature 485 (7400): 631-634. |
| Emp_25 | 89 | 125.75 | Brown, Joseph W., Robert B. Payne, David P. Mindell. 2007. Comment. Nuclear DNA does not reconcile ‘rocks’ and ‘clocks’ in Neoaves: a comment on Ericson et al. Biology Letters 3 (3): 257-259. |
| Emp_26 | 75 | 30.61 | Chris J. Law, Graham J. Slater, Rita S. Mehta, 2017, 'Lineage Diversity and Size Disparity in Musteloidea: Testing Patterns of Adaptive Radiation Using Molecular and Fossil-Based Methods', Systematic Biology |
| Emp_27 | 133 | 53.26 | Andersen, Michael J., Jenna M. McCullough, William M. Mauck, Brian Tilston Smith, Robert G. Moyle. 2017. A phylogeny of kingfishers reveals an Indomalayan origin and elevated rates of diversification on oceanic islands. Journal of Biogeography |

Table S1 Continued

| TreeID | Taxa | Age | Source |
|---|---|---|---|
| Emp_28 | 68 | 0.57 | Ronquist, Fredrik, Seraina Klopfstein, Lars Vilhelmsen, Susanne Schulmeister, Debra L. Murray, Alexandr P. Rasnitsyn. 2012. A Total-evidence approach to dating with fossils, applied to the early radiation of the Hymenoptera. Systematic Biology, 61(6): 973-999. |
| Emp_29 | 68 | 0.27 | Ronquist, Fredrik, Seraina Klopfstein, Lars Vilhelmsen, Susanne Schulmeister, Debra L. Murray, Alexandr P. Rasnitsyn. 2012. A Total-evidence approach to dating with fossils, applied to the early radiation of the Hymenoptera. Systematic Biology, 61(6): 973-999. |
| Emp_30 | 68 | 350.56 | Ronquist, Fredrik, Seraina Klopfstein, Lars Vilhelmsen, Susanne Schulmeister, Debra L. Murray, Alexandr P. Rasnitsyn. 2012. A Total-evidence approach to dating with fossils, applied to the early radiation of the Hymenoptera. Systematic Biology, 61(6): 973-999. |
| Emp_31 | 68 | 388.88 | Ronquist, Fredrik, Seraina Klopfstein, Lars Vilhelmsen, Susanne Schulmeister, Debra L. Murray, Alexandr P. Rasnitsyn. 2012. A Total-evidence approach to dating with fossils, applied to the early radiation of the Hymenoptera. Systematic Biology, 61(6): 973-999. |
| Emp_32 | 68 | 0.33 | Ronquist, Fredrik, Seraina Klopfstein, Lars Vilhelmsen, Susanne Schulmeister, Debra L. Murray, Alexandr P. Rasnitsyn. 2012. A Total-evidence approach to dating with fossils, applied to the early radiation of the Hymenoptera. Systematic Biology, 61(6): 973-999. |
| Emp_33 | 68 | 0.37 | Ronquist, Fredrik, Seraina Klopfstein, Lars Vilhelmsen, Susanne Schulmeister, Debra L. Murray, Alexandr P. Rasnitsyn. 2012. A Total-evidence approach to dating with fossils, applied to the early radiation of the Hymenoptera. Systematic Biology, 61(6): 973-999. |
| Emp_34 | 48 | 101.64 | Jarvis, E. D., S. Mirarab, A. J. Aberer, B. Li, P. Houde, C. Li, S. Y. W. Ho, B. C. Faircloth, B. Nabholz, J. T. Howard, A. Suh, C. C. Weber, R. R. da Fonseca, J. Li, F. Zhang, H. Li, L. Zhou, N. Narula, L. Liu, G. Ganapathy, B. Boussau, M. S. Bayzid, V. Zavidovych, S. Subramanian, T. Gabaldon, S. Capella-Gutierrez, J. Huerta-Cepas, B. Rekepalli, K. Munch, M. Schierup, B. Lindow, W. C. Warren, D. Ray, R. E. Green, M. W. Bruford, X. Zhan, A. Dixon, S. Li, N. Li, Y. Huang, E. P. Derryberry, M. F. Bertelsen, F. H. Sheldon, R. T. Brumfield, C. V. Mello, P. V. Lovell, M. Wirthlin, M. P. C. Schneider, F. Prosdocimi, J. A. Samaniego, A. M. V. Velazquez, A. Alfaro-Nunez, P. F. Campos, B. Petersen, T. Sicheritz-Ponten, A. Pas, T. Bailey, P. Scofield, M. Bunce, D. M. Lambert, Q. Zhou, P. Perelman, A. C. Driskell, B. Shapiro, Z. Xiong, Y. Zeng, S. Liu, Z. Li, B. Liu, K. Wu, J. Xiao, X. Yinqi, Q. Zheng, Y. Zhang, H. Yang, J. Wang, L. Smeds, F. E. Rheindt, M. Braun, J. Fjeldsa, L. Orlando, F. K. Barker, K. A. Jonsson, W. Johnson, K.-P. Koepfli, S. O'Brien, D. Haussler, O. A. Ryder, C. Rahbek, E. Willerslev, G. R. Graves, T. C. Glenn, J. McCormack, D. Burt, H. Ellegren, P. Alstrom, S. V. Edwards, A. Stamatakis, D. P. Mindell, J. Cracraft, E. L. Braun, T. Warnow, W. Jun, M. T. P. Gilbert, G. Zhang. 2014. Whole-genome analyses resolve early branches in the tree of life of modern birds. Science 346 (6215): 1320-1331. |
| Emp_35 | 95 | 97.11 | Dahiana Arcila, R. Alexander Pyron, James C. Tyler, Guillermo Orti, Ricardo Betancur-R., 2015, 'An evaluation of fossil tip-dating versus node-age calibrations in tetraodontiform fishes (Teleostei: Percomorphaceae)', Molecular Phylogenetics and Evolution, vol. 82, pp. 131-145 |
| Emp_36 | 43 | 390.58 | Dornburg, Alex, Jeffrey P Townsend, Matt Friedman, Thomas J Near. 2014. Phylogenetic informativeness reconciles ray-finned fish molecular divergence times. BMC Evolutionary Biology 14(1): 169. |

Table S1 Continued

| TreeID | Taxa | Age | Source |
|---|---|---|---|
| Emp_37 | 43 | 394.30 | Dornburg, Alex, Jeffrey P Townsend, Matt Friedman, Thomas J Near. 2014. Phylogenetic informativeness reconciles ray-finned fish molecular divergence times. BMC Evolutionary Biology 14(1): 169. |
| Emp_38 | 45 | 26.84 | Schweizer, Manuel, Timothy F. Wright, Joshua V. Penalba, Erin E. Schirtzinger, Leo Joseph. 2015. Molecular phylogenetics suggests a New Guinean origin and frequent episodes of founder-event speciation in the nectarivorous lories and lorikeets (Aves: Psittaciformes). Molecular Phylogenetics and Evolution 90: 34-48. |
| Emp_39 | 83 | 138.30 | Sean G Brady, Brian L Fisher, Ted R Schultz, Philip S Ward, 2014, 'The rise of army ants and their relatives: diversification of specialized predatory doryline ants', BMC Evolutionary Biology, vol. 14, no. 1, p. 93 |
| Emp_40 | 32 | 86.54 | Garcia-R, Juan C., Gillian C. Gibb, Steve A. Trewick. 2014. Eocene diversification of crown group rails (Aves: Gruiformes: Rallidae). PLoS ONE 9 (10): e109635 |
| Emp_41 | 124 | 514.26 | Beaulieu, Jeremy M., Brian C. O'Meara, Peter Crane, Michael J. Donoghue. 2015. Heterogeneous rates of molecular evolution and diversification could explain the Triassic age estimate for Angiosperms. Systematic Biology 64 (5): 869-878 |
| Emp_42 | 82 | 48.25 | K. M. Kozak, N. Wahlberg, A. F. E. Neild, K. K. Dasmahapatra, J. Mallet, C. D. Jiggins, 2015, 'Multilocus Species Trees Show the Recent Adaptive Radiation of the Mimetic Heliconius Butterflies', Systematic Biology, vol. 64, no. 3, pp. 505-524 |
| Emp_43 | 198 | 78.35 | Prum, Richard O., Jacob S. Berv, Alex Dornburg, Daniel J. Field, Jeffrey P. Townsend, Emily Moriarty Lemmon, Alan R. Lemmon. 2015. A comprehensive phylogeny of birds (Aves) using targeted next-generation DNA sequencing. Nature 526, (7574): 569-573 |
| Emp_44 | 198 | 72.91 | Prum, Richard O., Jacob S. Berv, Alex Dornburg, Daniel J. Field, Jeffrey P. Townsend, Emily Moriarty Lemmon, Alan R. Lemmon. 2015. A comprehensive phylogeny of birds (Aves) using targeted next-generation DNA sequencing. Nature 526, (7574): 569-573 |
| Emp_45 | 180 | 56.07 | Dufort, Matthew J. 2015. An augmented supermatrix phylogeny of the avian family Picidae reveals uncertainty deep in the family tree. Molecular Phylogenetics and Evolution |
| Emp_46 | 54 | 750.70 | dos Reis, Mario, Yuttapong Thawornwattana, Konstantinos Angelis, Maximilian J. Telford, Philip C.J. Donoghue, Ziheng Yang. 2015. Uncertainty in the timing of origin of animals and the limits of precision in molecular timescales. Current Biology |
| Emp_47 | 54 | 750.76 | dos Reis, Mario, Yuttapong Thawornwattana, Konstantinos Angelis, Maximilian J. Telford, Philip C.J. Donoghue, Ziheng Yang. 2015. Uncertainty in the timing of origin of animals and the limits of precision in molecular timescales. Current Biology |
| Emp_48 | 54 | 819.73 | dos Reis, Mario, Yuttapong Thawornwattana, Konstantinos Angelis, Maximilian J. Telford, Philip C.J. Donoghue, Ziheng Yang. 2015. Uncertainty in the timing of origin of animals and the limits of precision in molecular timescales. Current Biology |
| Emp_49 | 54 | 819.58 | dos Reis, Mario, Yuttapong Thawornwattana, Konstantinos Angelis, Maximilian J. Telford, Philip C.J. Donoghue, Ziheng Yang. 2015. Uncertainty in the timing of origin of animals and the limits of precision in molecular timescales. Current Biology |

Table S1 Continued

| TreeID | Taxa | Age | Source |
|---|---|---|---|
| Emp_50 | 54 | 776.48 | dos Reis, Mario, Yuttapong Thawornwattana, Konstantinos Angelis, Maximilian J. Telford, Philip C.J. Donoghue, Ziheng Yang. 2015. Uncertainty in the timing of origin of animals and the limits of precision in molecular timescales. Current Biology |
| Emp_51 | 54 | 776.34 | dos Reis, Mario, Yuttapong Thawornwattana, Konstantinos Angelis, Maximilian J. Telford, Philip C.J. Donoghue, Ziheng Yang. 2015. Uncertainty in the timing of origin of animals and the limits of precision in molecular timescales. Current Biology |
| Emp_52 | 54 | 813.44 | dos Reis, Mario, Yuttapong Thawornwattana, Konstantinos Angelis, Maximilian J. Telford, Philip C.J. Donoghue, Ziheng Yang. 2015. Uncertainty in the timing of origin of animals and the limits of precision in molecular timescales. Current Biology |
| Emp_53 | 54 | 813.69 | dos Reis, Mario, Yuttapong Thawornwattana, Konstantinos Angelis, Maximilian J. Telford, Philip C.J. Donoghue, Ziheng Yang. 2015. Uncertainty in the timing of origin of animals and the limits of precision in molecular timescales. Current Biology |
| Emp_54 | 88 | 68.80 | Cushing, Paula E., Matthew R. Graham, Lorenzo Prendini, Jack O. Brookhart. 2015. A multilocus molecular phylogeny of the endemic North American camel spider family Eremobatidae (Arachnida: Solifugae). Molecular Phylogenetics and Evolution 92: 280-293 |
| Emp_55 | 187 | 97.76 | Toussaint, Emmanuel F. A., Lars Hendrich, Helena Shaverdo, Michael Balke. 2015. Mosaic patterns of diversification dynamics following the colonization of Melanesian islands. Scientific Reports 5: 16016 |
| Emp_56 | 86 | 90.74 | McCord, Charlene L., Mark W. Westneat. 2016. Phylogenetic relationships and the evolution of BMP4 in triggerfishes and filefishes (Balistoidea). Molecular Phylogenetics and Evolution 94: 397-409 |
| Emp_57 | 48 | 96.57 | Claramunt, Santiago, Joel Cracraft. 2015. A new time tree reveals Earth historys imprint on the evolution of modern birds. Science Advances 1 (11): e1501005-e1501005 |
| Emp_58 | 102 | 62.58 | Gibb, Gillian C., Ryan England, Gerrit Hartig, P.A. (Trish) McLenachan, Briar L. Taylor Smith, Bennet J. McComish, Alan Cooper, David Penny. 2015. New Zealand passerines help clarify the diversification of major songbird lineages during the Oligocene. Genome Biology and Evolution 7 (11): 2983-2995. |
| Emp_59 | 72 | 81.00 | Ksepka, Daniel T., Matthew J. Phillips. 2015. Avian diversification patterns across the K-Pg boundary: influence of calibrations, datasets, and model misspecification. Annals of the Missouri Botanical Garden 100 (4): 300-328 |
| Emp_60 | 72 | 81.00 | Ksepka, Daniel T., Matthew J. Phillips. 2015. Avian diversification patterns across the K-Pg boundary: influence of calibrations, datasets, and model misspecification. Annals of the Missouri Botanical Garden 100 (4): 300-328 |
| Emp_61 | 72 | 104.07 | Ksepka, Daniel T., Matthew J. Phillips. 2015. Avian diversification patterns across the K-Pg boundary: influence of calibrations, datasets, and model misspecification. Annals of the Missouri Botanical Garden 100 (4): 300-328 |
| Emp_62 | 72 | 97.63 | Ksepka, Daniel T., Matthew J. Phillips. 2015. Avian diversification patterns across the K-Pg boundary: influence of calibrations, datasets, and model misspecification. Annals of the Missouri Botanical Garden 100 (4): 300-328 |
| Emp_63 | 72 | 111.90 | Ksepka, Daniel T., Matthew J. Phillips. 2015. Avian diversification patterns across the K-Pg boundary: influence of calibrations, datasets, and model misspecification. Annals of the Missouri Botanical Garden 100 (4): 300-328 |

Table S1 Continued

| TreeID | Taxa | Age | Source |
|---|---|---|---|
| Emp_64 | 72 | 111.07 | Ksepka, Daniel T., Matthew J. Phillips. 2015. Avian diversification patterns across the K-Pg boundary: influence of calibrations, datasets, and model misspecification. Annals of the Missouri Botanical Garden 100 (4): 300-328 |
| Emp_65 | 72 | 112.80 | Ksepka, Daniel T., Matthew J. Phillips. 2015. Avian diversification patterns across the K-Pg boundary: influence of calibrations, datasets, and model misspecification. Annals of the Missouri Botanical Garden 100 (4): 300-328 |
| Emp_66 | 72 | 85.96 | Ksepka, Daniel T., Matthew J. Phillips. 2015. Avian diversification patterns across the K-Pg boundary: influence of calibrations, datasets, and model misspecification. Annals of the Missouri Botanical Garden 100 (4): 300-328 |
| Emp_67 | 72 | 103.89 | Ksepka, Daniel T., Matthew J. Phillips. 2015. Avian diversification patterns across the K-Pg boundary: influence of calibrations, datasets, and model misspecification. Annals of the Missouri Botanical Garden 100 (4): 300-328 |
| Emp_68 | 72 | 81.35 | Ksepka, Daniel T., Matthew J. Phillips. 2015. Avian diversification patterns across the K-Pg boundary: influence of calibrations, datasets, and model misspecification. Annals of the Missouri Botanical Garden 100 (4): 300-328 |
| Emp_69 | 72 | 80.99 | Ksepka, Daniel T., Matthew J. Phillips. 2015. Avian diversification patterns across the K-Pg boundary: influence of calibrations, datasets, and model misspecification. Annals of the Missouri Botanical Garden 100 (4): 300-328 |
| Emp_70 | 72 | 106.29 | Ksepka, Daniel T., Matthew J. Phillips. 2015. Avian diversification patterns across the K-Pg boundary: influence of calibrations, datasets, and model misspecification. Annals of the Missouri Botanical Garden 100 (4): 300-328 |
| Emp_71 | 72 | 94.14 | Ksepka, Daniel T., Matthew J. Phillips. 2015. Avian diversification patterns across the K-Pg boundary: influence of calibrations, datasets, and model misspecification. Annals of the Missouri Botanical Garden 100 (4): 300-328 |
| Emp_72 | 72 | 81.08 | Ksepka, Daniel T., Matthew J. Phillips. 2015. Avian diversification patterns across the K-Pg boundary: influence of calibrations, datasets, and model misspecification. Annals of the Missouri Botanical Garden 100 (4): 300-328 |
| Emp_73 | 72 | 81.31 | Ksepka, Daniel T., Matthew J. Phillips. 2015. Avian diversification patterns across the K-Pg boundary: influence of calibrations, datasets, and model misspecification. Annals of the Missouri Botanical Garden 100 (4): 300-328 |
| Emp_74 | 72 | 81.01 | Ksepka, Daniel T., Matthew J. Phillips. 2015. Avian diversification patterns across the K-Pg boundary: influence of calibrations, datasets, and model misspecification. Annals of the Missouri Botanical Garden 100 (4): 300-328 |
| Emp_75 | 72 | 80.83 | Ksepka, Daniel T., Matthew J. Phillips. 2015. Avian diversification patterns across the K-Pg boundary: influence of calibrations, datasets, and model misspecification. Annals of the Missouri Botanical Garden 100 (4): 300-328 |
| Emp_76 | 35 | 104.99 | Delsuc, Frederic, Gillian C. Gibb, Melanie Kuch, Guillaume Billet, Lionel Hautier, John Southon, Jean-Marie Rouillard, Juan Carlos Fernicola, Sergio F. Vizcaino, Ross D.E. MacPhee, Hendrik N. Poinar. 2016. The phylogenetic affinities of the extinct glyptodonts. Current Biology 26 (4): R155-R156 |
| Emp_77 | 40 | 102.78 | Gibb, Gillian C., Fabien L. Condamine, Melanie Kuch, Jacob Enk, Nadia Moraes-Barros, Mariella Superina, Hendrik N. Poinar, Frederic Delsuc. 2015. Shotgun mitogenomics provides a reference phylogenetic framework and timescale for living Xenarthrans. Molecular Biology and Evolution 33 (3): 621-642 |
| Emp_78 | 100 | 78.41 | A. C. Schneider, W. A. Freyman, C. M. Guilliams, Y. P. Springer, B. G. Baldwin, 2016, 'Pleistocene radiation of the serpentine-adapted genus Hesperolinon and other divergence times in Linaceae (Malpighiales)', American Journal of Botany, vol. 103, no. 2, pp. 221-232 |

Table S1 Continued

| TreeID | Taxa | Age | Source |
|---|---|---|---|
| Emp_79 | 32 | 4.37 | Joel P. Olfelt, William A. Freyman, 2014, ' Relationships of North American members of Rhodiola (Crassulaceae) ', Botany, vol. 92, no. 12, pp. 901-910 |
| Emp_80 | 94 | 54.58 | Zhi-Yong Yuan, Wei-Wei Zhou, Xin Chen, Nikolay A. Poyarkov, Hong-Man Chen, Nian-Hong Jang-Liaw, Wen-Hao Chou, Nicholas J. Matzke, Koji Iizuka, Mi-Sook Min, Sergius L. Kuzmin, Ya-Ping Zhang, David C. Cannatella, David M. Hillis, Jing Che, 2016, 'Spatiotemporal Diversification of the True Frogs (Genus Rana): A Historical Framework for a Widely Studied Group of Model Organisms', Systematic Biology, p. syw055 |
| Emp_81 | 93 | 83.32 | Wang, Ning, Rebecca T. Kimball, Edward L. Braun, Bin Liang, Zhengwang Zhang. 2016. Ancestral range reconstruction of Galliformes: the effects of topology and taxon sampling. Journal of Biogeography |
| Emp_82 | 55 | 73.64 | Ericson, Per GP, Seraina Klopfstein, Martin Irestedt, Jacqueline MT Nguyen and Johan AA Nylander. 2014. Dating the diversification of the major lineages of Passeriformes (Aves). BMC Evolutionary Biology, 14(8). |
| Emp_83 | 34 | 0.53 | Slager, David L., C.J. Battey, Robert W. Bryson, Gary Voelker, John Klicka. 2014. A multilocus phylogeny of a major New World avian radiation: The Vireonidae. Molecular Phylogenetics and Evolution 80: 95-104 |
| Emp_84 | 34 | 0.60 | Slager, David L., C.J. Battey, Robert W. Bryson, Gary Voelker, John Klicka. 2014. A multilocus phylogeny of a major New World avian radiation: The Vireonidae. Molecular Phylogenetics and Evolution 80: 95-104 |
| Emp_85 | 36 | 21.90 | Sweet, Andrew D., Kevin P. Johnson. 2015. Patterns of diversification in small New World ground doves are consistent with major geologic events. The Auk 132 (1): 300-312. |
| Emp_86 | 76 | 223.77 | Martin Malmstrom, Michael Matschiner, Ole K Torresen, Bastiaan Star, Lars G Snipen, Thomas F Hansen, Helle T Baalsrud, Alexander J Nederbragt, Reinhold Hanel, Walter Salzburger, Nils C Stenseth, Kjetill S Jakobsen, Sissel Jentoft, 2016, 'Evolution of the immune system influences speciation rates in teleost fishes', Nature Genetics |
| Emp_87 | 64 | 0.17 | Wood, Jamie R., Kieren J. Mitchell, R. Paul Scofield, Vanesa L. De Pietri, Nicolas J. Rawlence, Alan Cooper. 2016. Phylogenetic relationships and terrestrial adaptations of the extinct laughing owl, Sceloglaux albifacies (Aves: Strigidae). Zoological Journal of the Linnean Society |
| Emp_88 | 106 | 4.43 | Moyle, Robert G., Carl H. Oliveros, Michael J. Andersen, Peter A. Hosner, Brett W. Benz, Joseph D. Manthey, Scott L. Travers, Rafe M. Brown, Brant C. Faircloth. 2016. Tectonic collision and uplift of Wallacea triggered the global songbird radiation. Nature Communications 7: 12709 |
| Emp_89 | 51 | 2.43 | Fuchs, Jerome, Jan I. Ohlson, Per G. P. Ericson, Eric Pasquet. 2007. Synchronous intercontinental splits between assemblages of woodpeckers suggested by molecular data. Zoologica Scripta 36 (1): 11-25 |
| Emp_90 | 33 | 126.66 | Johnson, Jeff A., Joseph W. Brown, Jerome Fuchs, David P. Mindell, 2016, 'Multi-locus phylogenetic inference among New World Vultures (Aves: Cathartidae)', Molecular Phylogenetics and Evolution, vol. 105, pp. 193-199 |
| Emp_91 | 33 | 126.76 | Johnson, Jeff A., Joseph W. Brown, Jerome Fuchs, David P. Mindell, 2016, 'Multi-locus phylogenetic inference among New World Vultures (Aves: Cathartidae)', Molecular Phylogenetics and Evolution, vol. 105, pp. 193-199 |
| Emp_92 | 195 | 339.35 | Foster, Charles S. P., Herve Sauquet, Marlien van der Merwe, Hannah McPherson, Maurizio Rossetto, Simon Y. W. Ho. 2016. Evaluating the impact of genomic data and priors on Bayesian estimates of the angiosperm evolutionary timescale. Systematic Biology, p. 66(3) |

Table S1 Continued

| TreeID | Taxa | Age | Source |
|---|---|---|---|
| Emp_93 | 107 | 17.11 | Scofield, R. Paul, Kieren J. Mitchell, Jamie R. Wood, Vanesa L. De Pietri, Scott Jarvie, Bastien Llamas, Alan Cooper. 2016. The Origin and Phylogenetic Relationships of the New Zealand Ravens. Molecular Phylogenetics and Evolution 106: 136-143 |
| Emp_94 | 84 | 65.00 | Jennifer L. Fessler, Mark W. Westneat, 2007, 'Molecular phylogenetics of the butterflyfishes (Chaetodontidae): Taxonomy and biogeography of a global coral reef fish family', Molecular Phylogenetics and Evolution, vol. 45, no. 1, pp. 50-68 |
| Emp_95 | 55 | 151.42 | Harrington, Richard C., Brant C. Faircloth, Ron I. Eytan, W. Leo Smith, Thomas J. Near, Michael E. Alfaro, Matt Friedman. 2016. Phylogenomic analysis of carangimorph fishes reveals flatfish asymmetry arose in a blink of the evolutionary eye. BMC Evolutionary Biology 16: 224 |
| Emp_96 | 108 | 56.36 | Lohmann, L. G., Bell, C. D., Calio, M. F., & Winkworth, R. C. (2013). Pattern and timing of biogeographical history in the Neotropical tribe Bignonieae (Bignoniaceae). Botanical Journal of the Linnean Society, 171(1), 154-170. |
| Emp_97 | 36 | 43.40 | Higdon J. W., Bininda-Emonds O. R. P., Beck R. M. D., Ferguson S. H. 2007. Phylogeny and divergence of the pinnipeds (Carnivora: Mammalia) assessed using a multigene dataset. BMC Evolutionary Biology 8: 216. |
| Emp_98 | 36 | 43.40 | Higdon J. W., Bininda-Emonds O. R. P., Beck R. M. D., Ferguson S. H. 2007. Phylogeny and divergence of the pinnipeds (Carnivora: Mammalia) assessed using a multigene dataset. BMC Evolutionary Biology 8: 216. |
| Emp_99 | 36 | 43.40 | Higdon J. W., Bininda-Emonds O. R. P., Beck R. M. D., Ferguson S. H. 2007. Phylogeny and divergence of the pinnipeds (Carnivora: Mammalia) assessed using a multigene dataset. BMC Evolutionary Biology 8: 216. |
| Emp_100 | 73 | 144.34 | Uit de weerd D.R., & Gittenberger E. 2013. Phylogeny of the land snail family Clausiliidae (Gastropoda: Pulmonata). Molecular Phylogenetics and Evolution, 67(1): 201-216. |
| Emp_101 | 157 | 48.96 | Hubert N., Paradis E., Bruggemann H., & Planes S. 2011. Community assembly and diversification in Indo-Pacific coral reef fishes. Ecology and Evolution, 1(3): 229-277. |
| Emp_102 | 37 | 0.17 | de Sa R.O., Streicher J.W., Selenkoyela R., Forlani M.C., Loader S.P., Greenbaum E., Richards S., & Haddad C. 2012. Molecular phylogeny of microhylid frogs (Anura: Microhylidae) with emphasis on relationships among New World genera. BMC Evolutionary Biology 12: 241 . |
| Emp_103 | 68 | 39.92 | Valtuena, Francisco J., Chris D. Preston, and Joachim W. Kadereit. "Phylogeography of a Tertiary relict plant, Meconopsis cambrica (Papaveraceae), implies the existence of northern refugia for a temperate herb." Molecular Ecology 21.6 (2012): 1423-1437. |
| Emp_104 | 87 | 35.86 | Steeman, M., Hebsgaard M., Fordyce R., Ho S., Rabosky D., Nielsen R., Rahbek C., Glenner H., Sorensen M., & Willerslev E. 2009. Radiation of Extant Cetaceans Driven by Restructuring of the Oceans. Systematic Biology 58 (6): 573-585. |
| Emp_105 | 171 | 49.38 | Lack J.B., & Van den bussche R.A. 2010. Identifying the Confounding Factors in Resolving Phylogenetic Relationships in Vespertilionidae. Journal of Mammalogy, . |
| Emp_106 | 154 | 41.92 | Dumont E.R., Davalos L.M., Goldberg A., Santana S.E., Rex K., & Voigt C.C. 2012. Morphological innovation, diversification and invasion of a new adaptive zone. Proceedings of the Royal Society B: Biological Sciences, 279: 1797-1805. |
| Emp_107 | 32 | 28.07 | Chan, L.M., Choi D., Raselimanana A.P., Rakotondravony H.E., & Yoder A.D. 2012. Defining spatial and temporal patterns of phylogeographic structure in Madagascar's iguanid lizards (genus Oplurus). Molecular Ecology 21 (15): 3839-3851. |

Table S1 Continued

| TreeID | Taxa | Age | Source |
|---|---|---|---|
| Emp_108 | 34 | 76.40 | Andujar, C., Serrano J., & Gomez-zurita J. 2012. Winding up the molecular clock in the genus Carabus (Coleoptera: Carabidae): assessment of methodological decisions on rate and node age estimation. BMC Evolutionary Biology 12: 40. |
| Emp_109 | 144 | 395.07 | K. Mao, R. I. Milne, L. Zhang, Y. Peng, J. Liu, P. Thomas, R. R. Mill, S. S. Renner. 2012. Distribution of living Cupressaceae reflects the breakup of Pangea. Proceedings of the National Academy of Sciences 109(20):7793-7798. |
| Emp_110 | 191 | 21.00 | Barker, F. K., K. J. Burns, J. Klicka, S. M. Lanyon, I. J. Lovette. 2013. Going to extremes: contrasting rates of diversification in a recent radiation of New World passerine birds. Systematic Biology 62 (2): 298-320. |
| Emp_111 | 115 | 100.00 | Lovette, Irby J., Jorge L. Perez-Eman, John P. Sullivan, Richard C. Banks, Isabella Fiorentino, Sergio Cordoba-Cordoba, Maria Echeverry-Galvis, F. Keith Barker, Kevin J. Burns, John Klicka, Scott M. Lanyon, Eldredge Bermingham. 2010. A comprehensive multilocus phylogeny for the wood-warblers and a revised classification of the Parulidae (Aves). Molecular Phylogenetics and Evolution 57 (2): 753-770. |
| Emp_112 | 63 | 187.94 | Sikes, D.S., & Venables C. 2013. Molecular phylogeny of the burying beetles (Coleoptera: Silphidae: Nicrophorinae). Molecular Phylogenetics and Evolution 69 (3): 552-565. |
| Emp_113 | 48 | 90.84 | Gibb, Gillian C., Martyn Kennedy, David Penny. 2013. Beyond phylogeny: pelecaniform and ciconiiform birds, and long-term niche stability. Molecular Phylogentics and Evolution 68 (2): 229-238. |
| Emp_114 | 176 | 90.38 | Bremer, B., & Eriksson, O. 2009. Time tree of Rubiaceae: Phylogeny and dating the family, subfamilies and tribes. Internat. J. Plant Sci. 170: 766-793. |
| Emp_115 | 44 | 81.74 | Chakrabarty, Prosanta, Matthew P. Davis, W. Leo Smith, Zachary H. Baldwin, John S. Sparks. 2011. Is sexual selection driving diversification of the bioluminescent ponyfishes (Teleostei: Leiognathidae)? Molecular Ecology 20 (13): 2818-2834. |
| Emp_116 | 154 | 122.75 | Friedman, M., B. P. Keck, A. Dornburg, R. I. Eytan, C. H. Martin, C. D. Hulsey, P. C. Wainwright, T. J. Near. 2013. Molecular and fossil evidence place the origin of cichlid fishes long after Gondwanan rifting. Proceedings of the Royal Society B: Biological Sciences 280 (1770): 20131733 |
| Emp_117 | 95 | 349.79 | Chen, Wei-Jen, Sebastien Lavoue, and Richard L. Mayden. 2013. Evolutionary origin and early biogeography of otophysan fishes (Ostariophysi: Teleostei). Evolution 67 (8): 2218-2239. |
| Emp_118 | 44 | 21.08 | Koepfli, Klaus-Peter, Kerry A Deere, Graham J Slater, Colleen Begg, Keith Begg, Lon Grassman, Mauro Lucherini, Geraldine Veron, Robert K Wayne. 2008. Multigene phylogeny of the Mustelidae: Resolving relationships, tempo and biogeographic history of a mammalian adaptive radiation. BMC Biology 6 (1): 10. |
| Emp_119 | 53 | 556.63 | Lee, M. S., Soubrier, J., & Edgecombe, G. D. (2013). Rates of phenotypic and genomic evolution during the Cambrian explosion. Current Biology 23 (19): 1889-1895. |
| Emp_120 | 137 | 16.64 | Drummond, C. S., Eastwood, R. J., Miotto, S. T., & Hughes, C. E. (2012). Multiple continental radiations and correlates of diversification in Lupinus (Leguminosae): testing for key innovation with incomplete taxon sampling. Systematic Biology 61(3), 443-460. |
| Emp_121 | 45 | 74.27 | Aggerbeck, Marie, Jon Fjeldsa, Les Christidis, Pierre-Henri Fabre, Knud Andreas Jonsson. 2014. Resolving deep lineage divergences in core corvoid passerine birds supports a proto-Papuan island origin. Molecular Phylogenetics and Evolution 70: 272-285. |

Table S1 Continued

| TreeID | Taxa | Age | Source |
|---|---|---|---|
| Emp_122 | 109 | 1781.09 | Parfrey, L. W., D. J. G. Lahr, A. H. Knoll, L. A. Katz. 2011. Estimating the timing of early eukaryotic diversification with multigene molecular clocks. Proceedings of the National Academy of Sciences 108 (33): 13624-13629. |
| Emp_123 | 93 | 54.58 | Smith, SA. 2009. Taking into account phylogenetic and divergence-time uncertainty in a parametric biogeographical analysis of the Northern Hemisphere plant clade Caprifolieae. Journal of Biogeography. |
| Emp_124 | 55 | 47.27 | Cibois, Alice, Jean-Claude Thibault, Caroline Bonillo, Christopher E. Filardi, Dick Watling, Eric Pasquet. 2014. Phylogeny and biogeography of the fruit doves (Aves: Columbidae). Molecular Phylogenetics and Evolution 70: 442-453. |
| Emp_125 | 44 | 45.48 | Drew, Bryan T., Kenneth J. Sytsma. 2013. The South American radiation of Lepechinia (Lamiaceae): phylogenetics, divergence times and evolution of dioecy. Botanical Journal of the Linnean Society 171 (1): 171-190. |
| Emp_126 | 100 | 50.00 | Mahler, D. L., T. Ingram, L. J. Revell, J. B. Losos. 2013. Exceptional Convergence on the Macroevolutionary Landscape in Island Lizard Radiations. Science 341 (6143): 292-295. |
| Emp_127 | 176 | 96.13 | Price, Trevor D., Daniel M. Hooper, Caitlyn D. Buchanan, Ulf S. Johansson, D. Thomas Tietze, Per Alstrom, Urban Olsson, Mousumi Ghosh-Harihar, Farah Ishtiaq, Sandeep K. Gupta, Jochen Martens, Bettina Harr, Pratap Singh, Dhananjai Mohan. 2014. Niche filling slows the diversification of Himalayan songbirds. Nature 509: 222-225. |
| Emp_128 | 34 | 244.52 | Mitchell, K. J., B. Llamas, J. Soubrier, N. J. Rawlence, T. H. Worthy, J. Wood, M. S. Y. Lee, A. Cooper. 2014. Ancient DNA reveals elephant birds and kiwi are sister taxa and clarifies ratite bird evolution. Science 344 (6186): 898-900. |
| Emp_129 | 108 | 92.98 | Antonelli, A. 2009. Have giant lobelias evolved several times independently? Life form shifts and historical biogeography of the cosmopolitan and highly diverse subfamily Loeblioideae (Campanulaceae). BMC Biology 7: 82 - see http://www.biomedcentral.com/1741-7007/7/82 |
| Emp_130 | 73 | 43.00 | Bergh N.G., & Linder H.P. 2009. Cape diversification and repeated out-of-southern-Africa dispersal in paper daisies. Molecular Phylogenetics and Evolution, 51(Special issue: origins and evolution of a biodiversity hotspot, the biodiversity of the African Cape Floristic Region): 5 - 18. |
| Emp_131 | 116 | 100.00 | Johnson, M. T., FitzJohn, R. G., Smith, S. D., Rausher, M. D., & Otto, S. P. (2011). Loss of sexual recombination and segregation is associated with increased diversification in evening primroses. Evolution, 65(11), 3230-3240. |
